# *ProteoSync*, a program for ortholog selection, automated sequence alignment and conservation projection onto protein atomic coordinates

**DOI:** 10.1101/2025.03.09.642228

**Authors:** Elliot Sicheri, Daniel Mao, Frank Sicheri

## Abstract

The projection of conservation onto the surface of a protein’s 3D structure is a powerful way of inferring functionally important regions. At present, the workflow for doing so can be involved and tedious. For this reason, we created ProteoSync, a Python program that semi-automates the process. The program creates an annotated sequence alignment of orthologs from a diverse set of selectable species, and enables the fast projection of amino acid conservation onto a predicted or known 3D model in PyMOL^[1]^.

## Introduction

At present, the workflow for identifying conserved and functionally important surfaces on a protein’s structure can be involved and tedious. First, one must manually search through candidate species for likely orthologs of the protein of interest and then use them to generate a multi protein sequence alignment to determine conservation. One must then manually project this conservation onto a protein’s structure, followed by visual inspection of the results. Since the process is iterative and requires inspection of the alignments to filter out non-orthologs (for example structural homologs), the tediousness of the process is amplified. For this reason, we created ProteoSync, a Python program that creates annotated sequence alignments of likely orthologs from a diverse, customizable set of species, and that enables the convenient projection of amino acid conservation scores onto a 3D model in PyMOL^[1]^. ProteoSync is available for download from GitHub (https://github.com/ElliotSicheri/ProteoSync) alongside a starter set of 84 diverse species sequence databases. The download is composed of a .zip file, which contains the executable, setup and usage instructions, database files, and subdirectories for outputs (**Figure 1**). A more detailed description of the downloaded contents is provided in the instructions folder.

**Figure 1.**
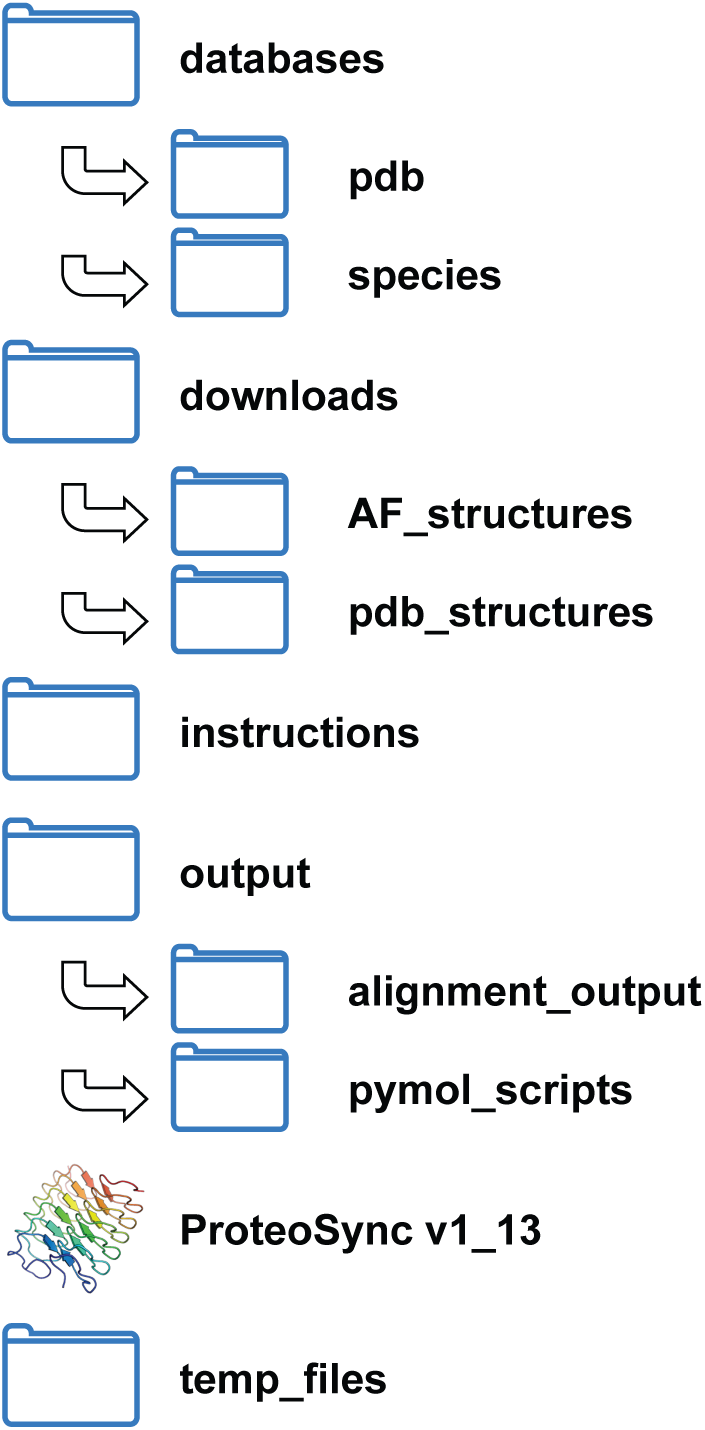
Directory structure of the ProteoSync installation folder.

## Methods

ProteoSync accepts a query protein sequence, a % identity threshold to filter likely functional orthologs, a list of species to analyze entered through a checkbox menu, optionally a UniProt ID ^[2]^ for the query protein of interest to enable the retrieval of an AlphaFold predicted structure ^[3]^ if available, and lastly an output file name. The program operates in 3 steps:

**Step 1:** Searching for protein orthologs: First the program uses the BLAST search algorithm^[4]^ to search a series of databases, each containing every protein sequence encoded in a particular species’ genome. Each species database provided in the software release was obtained from NCBI^[5]^ as described in **Addendum 1**. The set of species databases to be searched is defined by the user through a checkbox menu. In its downloaded configuration, databases for 84 searchable species are provided as listed in **Table 1**. However, the user can expand the set of species beyond the base set by following instructions described in **Addendum 1**.

**Table 1.**
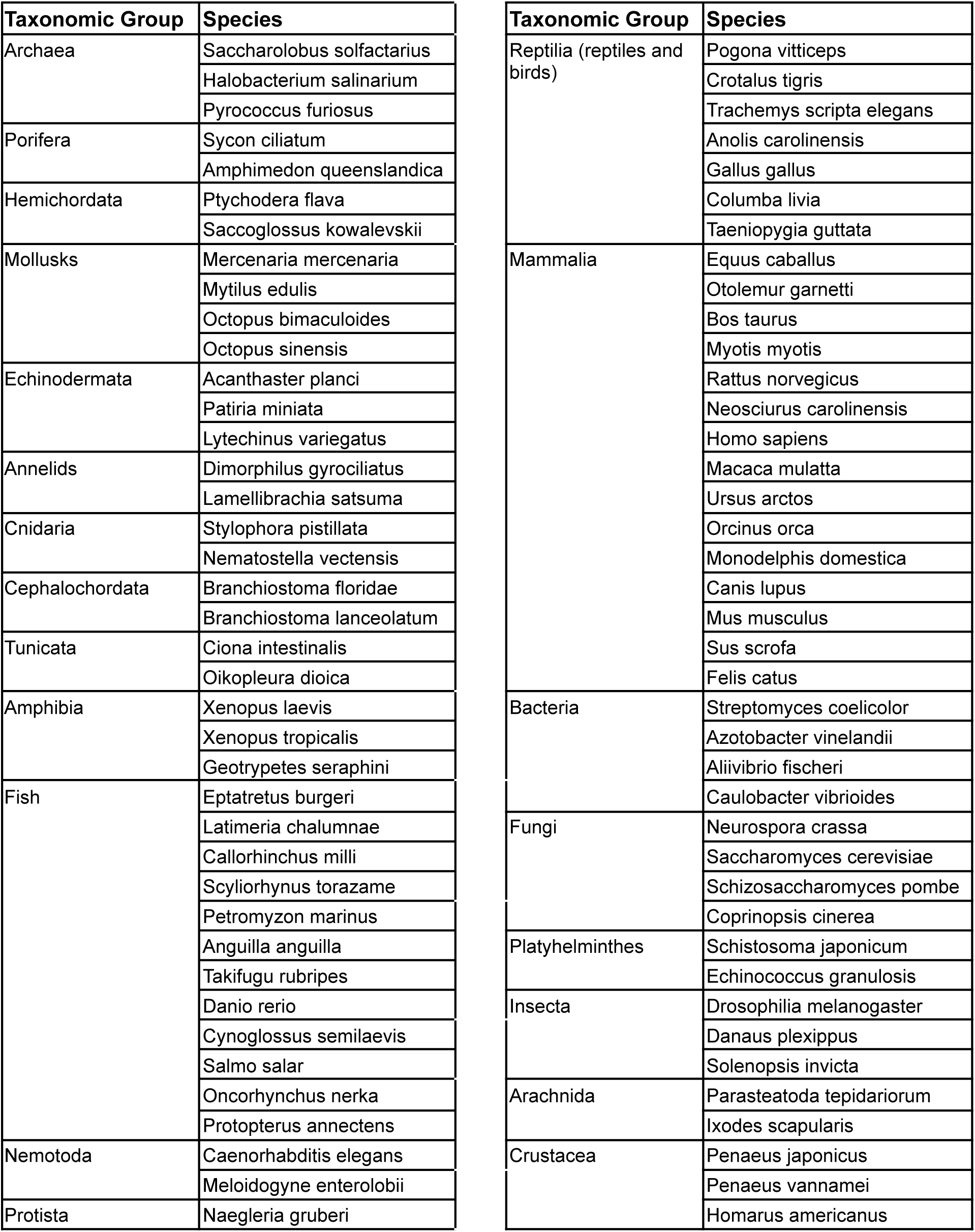
List of species available in the included starter species database set.

The BLAST search returns a single sequence most similar to the query sequence from each species database, which is most likely but not necessarily the direct functional ortholog to the query protein. As there is a possibility that the sequence identified is a protein homolog (same protein fold but divergent function) rather than a protein ortholog (same protein fold and same function), the user can define an identity threshold ranging from 0% to 100% to help filter out non-desirable protein sequences. All identified sequences with a % identity to the query sequence above the threshold are selected for sequence alignment. (**Figure 2**) Note that the identity threshold must be optimized for each query protein by iteratively running ProteoSync with different threshold settings until a desirable conservation projection is reached. However, a 50% cutoff threshold is a useful starting point. If the similarity threshold is set too high, too few sequences are selected, giving rise to too little evolutionary divergence, which results in the whole protein surface appearing as highly conserved. If the setting is set too low, structural homologs can be appropriately selected, which has the effect of masking or washing out the detection of conservation hotspots (i.e. the whole protein surface appears non-conserved).

**Figure 2.**
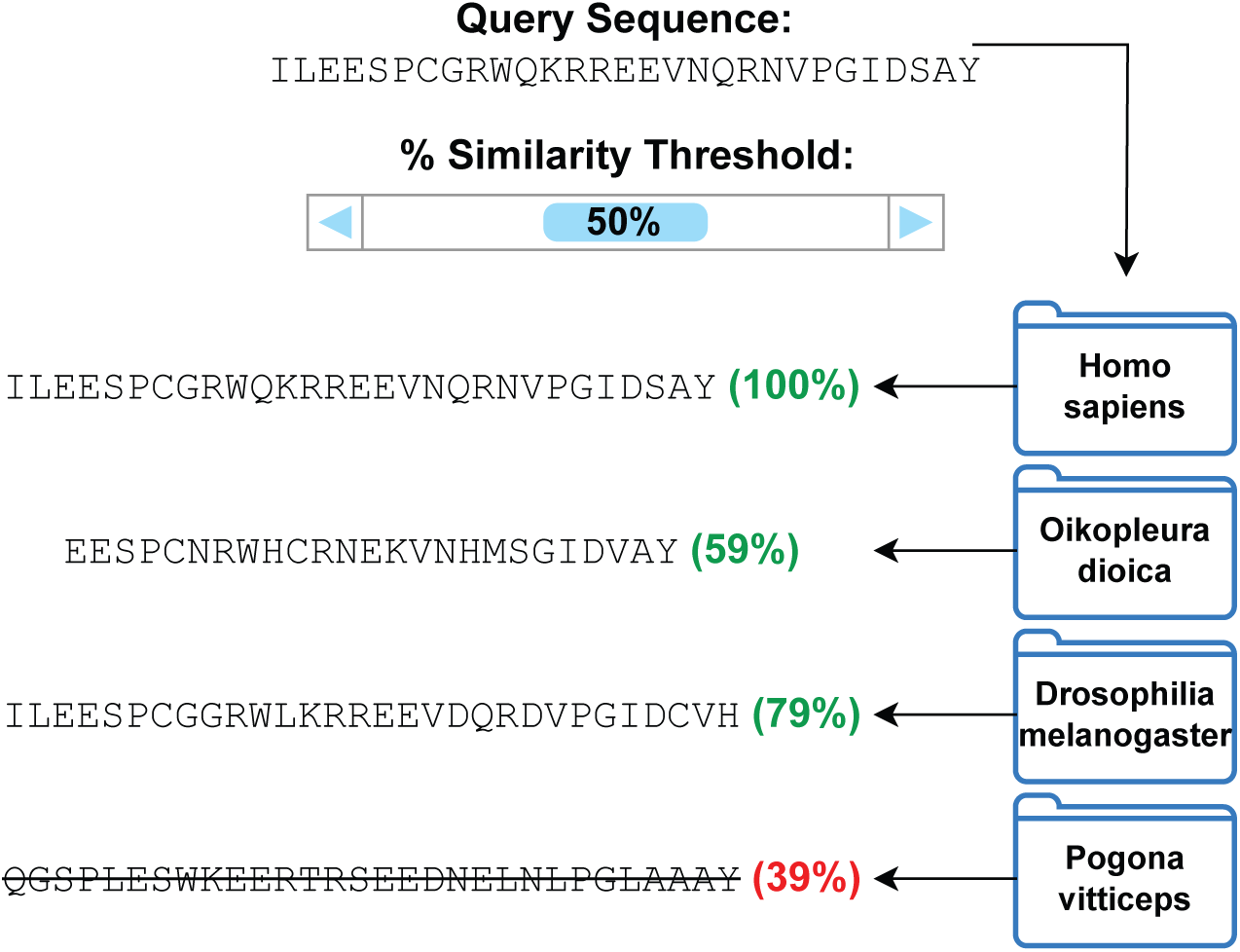
Example of the ortholog search process, employing a % identity threshold of 50%.

**Step 2:** Aligning sequences and determining conservation: ProteoSync then applies the clustalw sequence alignment algorithm^[6]^ to the predicted orthologous sequences to create a sequence alignment with conservation scores for each residue. The conservation scores are displayed (as in **Figure 3**) using common alignment score conventions^[7]^, where:

**Figure 3.**
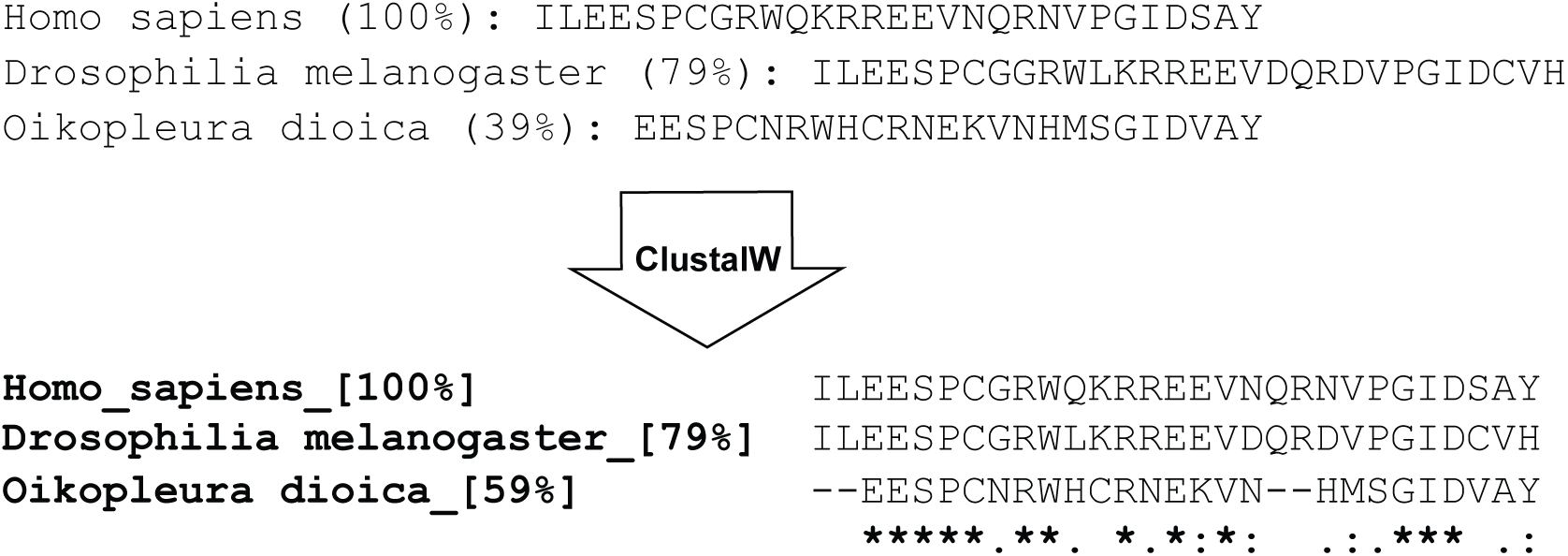
Example of output from the clustalw alignment algorithm.

- ‘*’ indicates that the residue is identical across all species.
- ‘:’ indicates that there are conservative substitutions, meaning the residues are very similar across species.
- ‘.’ indicates that there are semi-conservative substitutions, meaning the residues are fairly similar across the aligned species.
- ‘ ‘ indicates that there are non-conservative substitutions, meaning the residues are not similar between the aligned species.

**Step 3:** Secondary structure analysis: ProteoSync then uses the BLAST algorithm to search all protein sequences in the PDB (http://www.rcsb.org) for the closest sequence within a selectable identity threshold (same as that used for ortholog sequence search) for which an existing structure is available. A version of the PDB sequence database is provided in the software distribution package (ProteoSync/databases/PDB) and can be updated with the latest entries with the “update PDB database” button in the ProteoSync window. ProteoSync then downloads the PDB coordinate file (.pdb) for the best match into the ‘Proteosync/downloads/pdb_structures’ directory and extracts the positions of helices and beta sheets in the protein using the DSSP algorithm (Kabsch et al.)^[8]^. ProteoSync then aligns the secondary structure elements to the sequence alignment as a string of characters, where:

- ‘-’ indicates that the residue is unstructured
- ‘A’ indicates that the residue is part of an alpha helix
- ‘B’ indicates that the residue is part of a beta strand
- ‘X’ indicates that the residue is not modelled in the pdb structure

For large proteins, multiple structures in the PDB may be identified corresponding to individual domains or segments rather than one structure for a whole protein. If the top hit from the PDB sequence search excludes parts of the query sequence, the program runs iterative BLAST searches on the excluded regions to identify additional structure matches. All structures found are aligned on individual lines in the output alignment, labelled with their corresponding PDB codes. If the user entered a UniProt code^[2]^ for their protein, the program will also download the AlphaFold^[3]^ predicted structure for the query protein into the ‘Proteosync/downloads/af_structures’ directory, if available, and aligns its secondary structure to the sequence alignment in the same way as performed for the closest structure(s) in the PDB. The results are as displayed as in **Figure 4**. The final multi-protein sequence alignment with conservation and secondary structure projection is then outputted into a text file as shown in **Figure 5** in the ‘Proteosync/outputs/alignments’ directory. A list of accession codes for all sequences included in the alignment are included at the bottom of the file for easy referencing and validation.

**Figure 4.**
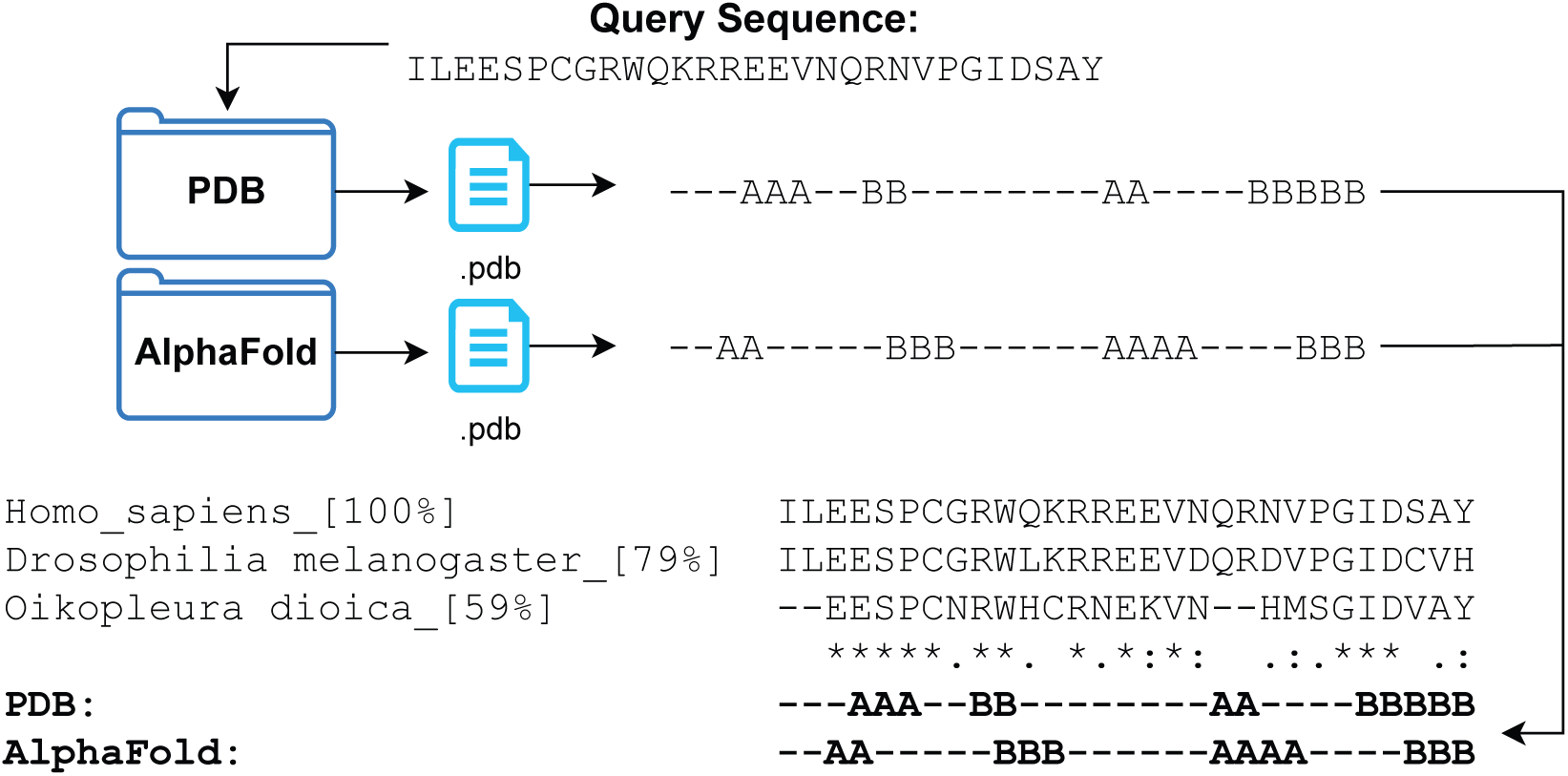
Example of output from the secondary structure alignment function using DSSP.

**Figure 5.**
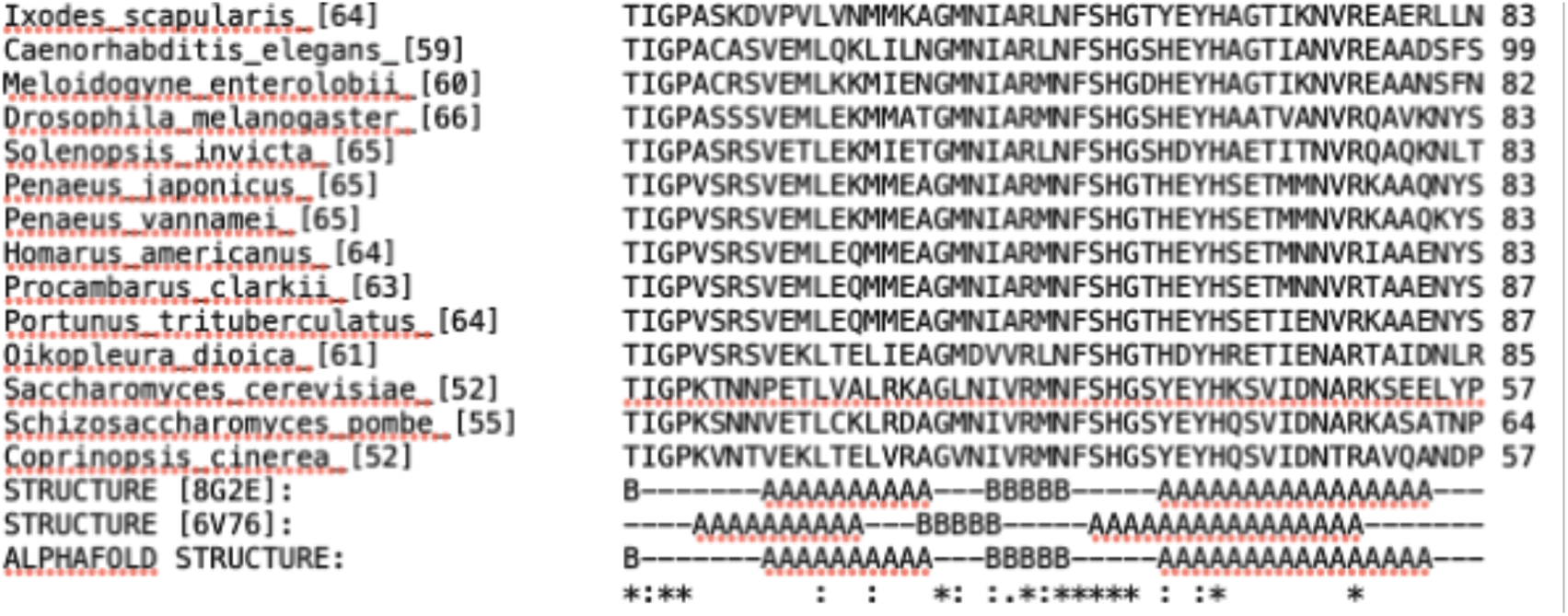
Example of a sequence alignment output file.

**Step 4.** Pymol Script Generation: Finally, a Pymol script file is created that allows projection of conservation from the alignment onto an empirically determined or predicted 3D model of the query protein downloaded by the user from the PDB (http://www.rcsb.org) or from the AlphaFold repository (https://alphafold.ebi.ac.uk/). This script file is stored in the

> ‘ProteoSync/outputs/pymol_scripts’ directory. The protein colouring convention corresponds to the clustalw alignment scores as follows:- Dark blue indicates that the residue is identical across all species (equivalent to ‘*’ in the alignment)

- Medium blue indicates that there are conservative substitutions, meaning the residues are very similar across species (equivalent to ‘:’ in the alignment)

- Light blue indicates that there are semi-conservative substitutions, meaning the residues are fairly similar across the aligned species. (equivalent to ‘.’ in the alignment)

- Grey indicates that there are non-conservative substitutions, meaning the residues are not similar between the aligned species. (equivalent to ‘ ‘ in the alignment)

## Results

To demonstrate the utility of the program, we ran ProteoSync on the human tumour suppressor protein FBW7 and the human pyruvate kinase PKM. FBW7 is an E3 ubiquitin ligase adapter subunit that is conserved from human to yeast (Cdc4 in *S. Cerevisiae*). It possesses a WD40-repeat domain that binds to conserved peptide elements termed degrons in target proteins, rendering them a substrate for protein ubiquitination and an F-box domain that enables its integration into a multi-subunit E3 ligase by binding to SKP1 (Hao et al 2007)^[9]^, (Orlicky et al.)^[10]^. ProteoSync analysis was performed using a 50% threshold cutoff on the entire starter set of 84 species and no intervention to remove faulty/truncated sequences from the resultant alignment (see **Supplementary Information Figure 1** for sequence alignment output). Projection of the calculated conservation scores onto the AlphaFold model of FBW7 revealed a hot spot of invariant conservation on the front face of the WD40-repeat domain, corresponding to the degron binding pocket (**Figure 6**).

**Figure 6.**
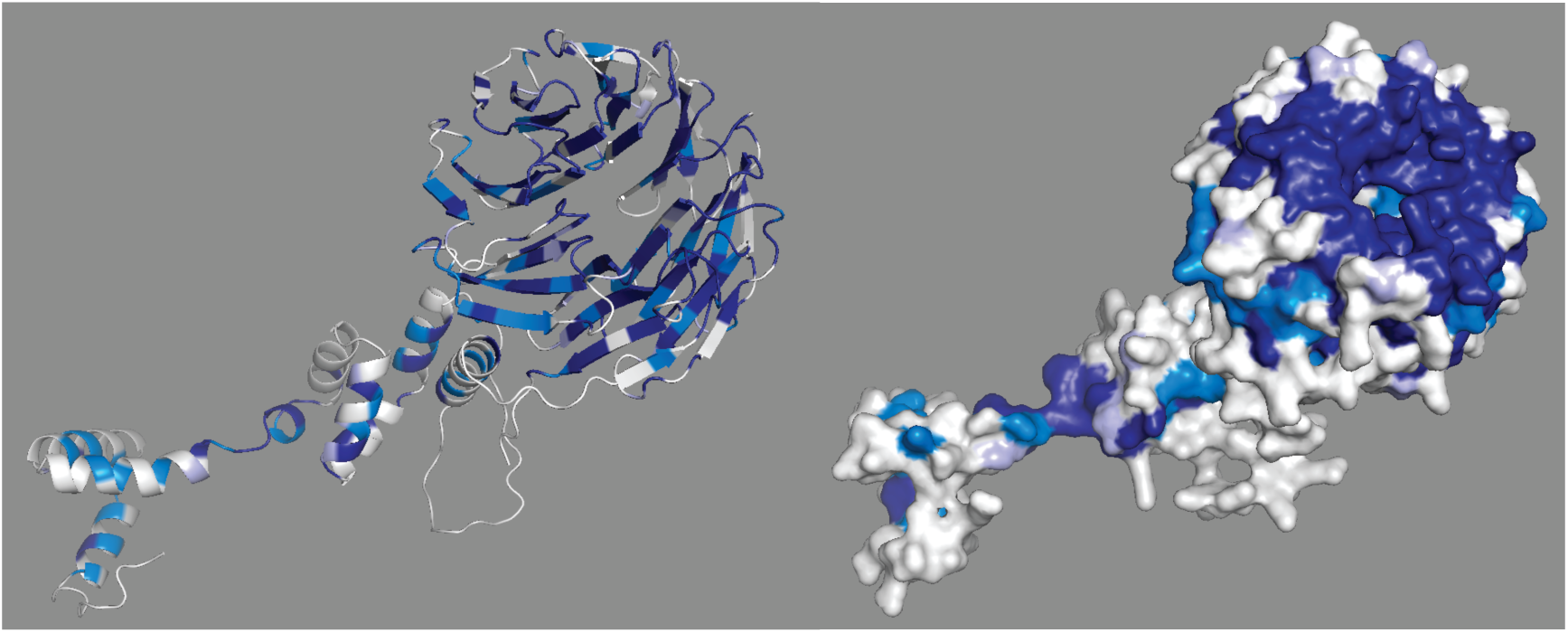
Conservation scores produced by ProteoSync projected onto the AlphaFold predicted structure of residues 222 to 707 of human FBW7. Shown are cartoon representation (left) and surface representation (right). Colouring is according to the ClustalW score generated in the output alignment (dark blue = invariant, medium blue = highly conserved (conservative substitutions), light blue = some conservation (semi-conservative substitutions), and white = not conserved). A hotspot of conservation on the front surface of the WD40 repeat domain identifies the functionally important degron binding pocket.

Human pyruvate kinase PKM is an enzyme responsible for catalyzing the final step in glycolysis, where the substrate phosphoenolpyruvate is converted into pyruvate, producing ATP (Dombrauckas et al. 2005)^[11]^. The active site of PKM houses the substrate binding and catalytic infrastructure in a cleft between the A domain (residues 44−116 and 219−389) and the B domain (residues 117-218) (Dombrauckas et al. 2005)^[11]^. ProteoSync analysis was performed using a 50% threshold cutoff on the entire starter species set with 7 species selected for exclusion due to the detection of faulty/truncated sequences in an alignment from an initial run of ProteoSync. (see **Supplementary Information Figure 2** for sequence alignment output). Projection of conservation onto an AlphaFold model of PKM revealed a hot spot of conservation along the cleft region between the two domains, while the rest of the protein was mostly unconserved (**Figure 7**).

**Figure 7.**
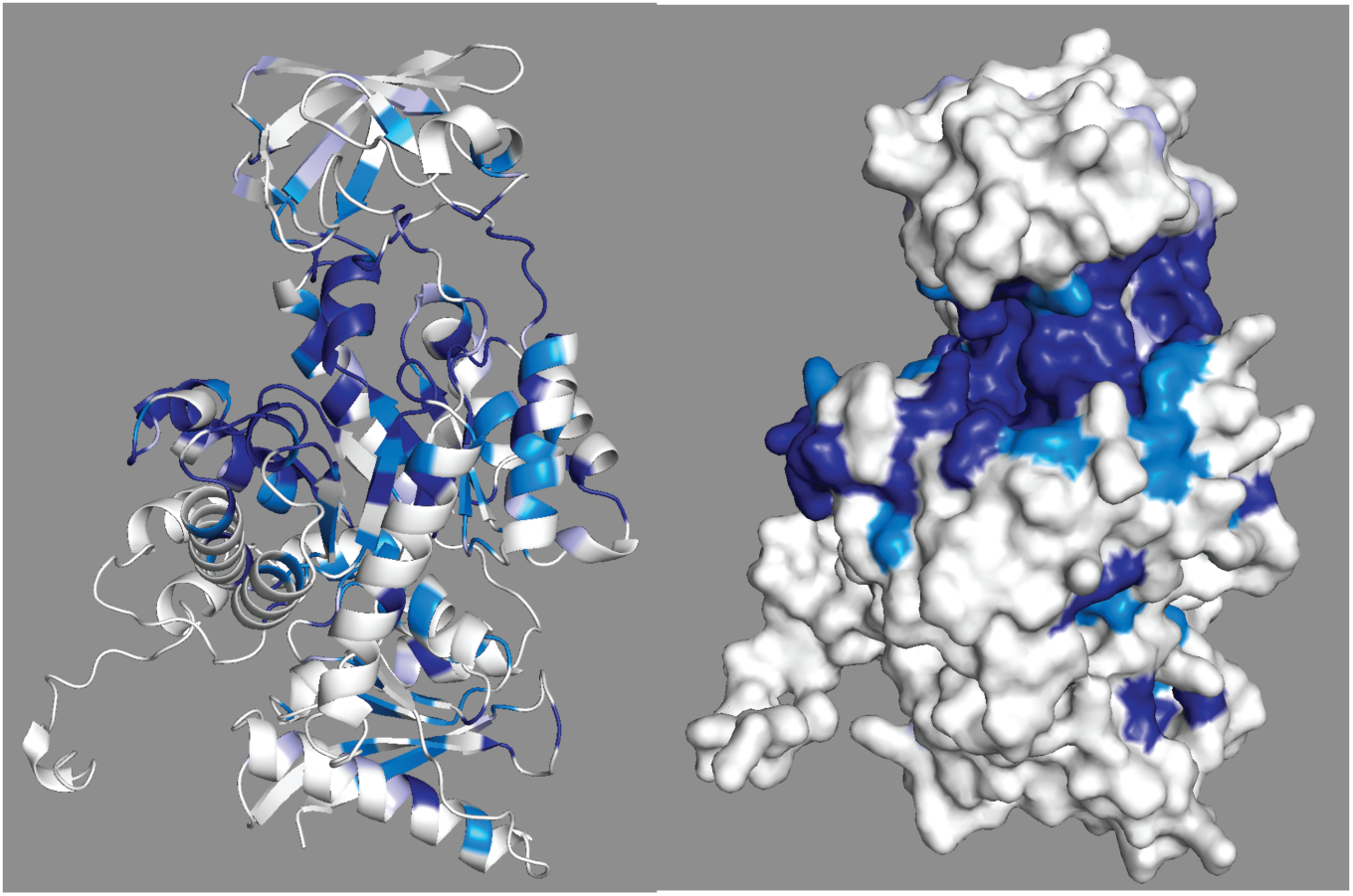
Conservation scores produced by ProteoSync projected onto the AlphaFold predicted structure of human PKM. Shown are cartoon representation (left) and surface representation (right). 7 species excluded from the starter species set due to faulty/truncated sequences include: Lamellibrachia satsuma, Mytilus edulis, Branchiostoma floridae, Branchiostoma lanceolatum, Columba livia, Otolemur garnettii, and Danaus plexippus. Colouring is according to the ClustalW score generated in the output alignment (dark blue = invariant, medium blue = highly conserved (conservative substitutions), light blue = some conservation (semi-conservative substitutions), and white = not conserved). A hotspot of conservation is evident in the cleft region between the upper and lower domains, corresponding to the functionally relevant active site.

## Discussion

In summary, ProteoSync is a program that can be used to quickly and easily calculate evolutionary conservation of a protein sequence across a user-defined set of species and project this conservation onto 3D models visualized by Pymol. Guided by visual inspection of the output, ProteoSync enables optimization of the clustalw alignment to obtain a more meaningful surface projection. Occasionally, one or more problematic sequences is encountered that lack a stretch of residues present in the bulk of other species sequences, potentially due to sequencing artifacts in the data obtained from NCBI. Due to the nature of the clustalW algorithm, these missing regions cause a complete loss of detail in the conservation scores, such that the region is indiscriminately marked as unconserved. Problematic sequences bearing these deletions are easily spotted in the clustalw output alignment file and can be removed via the taxonomy menu for the execution of a following ProteoSync cycle.

ProteoSync has many practical applications including assisting in the engineering of optimal protein constructs for crystallization and biochemical experimentation. Long, unstructured regions in a protein can inhibit protein crystallization and can lead to poor protein behavior in solution including aggregation and precipitation. While these unstructured segments can be readily identified in AlphaFold predicted models, conserved regions within flexible loops can contribute to protein folding or other conserved functions, which one may be keen to maintain. By projecting conservation generated by ProteoSync onto an AF-model, a better picture of what regions of a protein are likely dispensable or essential for a protein’s function can be achieved.

### Addendum 1. Instructions for installing ProteoSync dependencies

ProteoSync runs on all Apple computers tested. The program calls on several external programs which need to be installed separately, using the free program Anaconda. First, install Conda, available at https://www.anaconda.com/products/distribution. Note that irrespective of whether the computer employs an Apple silicon or Intel chipset, the user should install the Intel version of Anaconda, since some of the dependencies are only available for the Intel version.

Second, open a terminal window and install the following packages using the command lines:

- conda install -c salilab dssp

- conda install -c bioconda clustalw

- conda install -c bioconda blast

### Addendum 2. Instructions for synthesis of new species databases

Currently we have chosen a range of 84 species to include in our base installation, with a bias towards higher eukaryote diversity. There is a limited repertoire of bacterial, fungal and archaeal species. However, additional species can be added easily into the ProteoSync species dataset by following these instructions:

#### Step 1: Acquire a FASTA file containing all protein sequences of the species

A .fasta file containing all known sequences of a species’s genome can be obtained from NCBI by following these instructions. If you already have a .fasta file for a species of interest, you can skip this step.

- Get the species’ taxonomy id by searching for it at https://www.ncbi.nlm.nih.gov/Taxonomy/TaxIdentifier/tax_identifier.cgi
- On NCBI, select protein database and enter ‘txid####[Organism:noexp]’ (replace #### with the tax id)

- We recommend also adding ‘NOT partial’ to exclude partial sequences from the results
- Near the top left of the search results, click ‘Send to:’, select File, set Format to FASTA, and click Create File

#### Step 2: Format the FASTA file into a BLAST database

- To format the database, you’ll need the BLAST+ command line application. To install it, follow the instructions below.

- Go to https://ftp.ncbi.nlm.nih.gov/blast/executables/LATEST/
- Download ncbi-blast-2.14.0+.dmg
- Open the installer and follow the instructions
- Create a new folder with the name of your species as you want it to appear on the taxonomy menu.

- All spaces must be replaced by ‘_’.
- Try not to use any other special characters.
- Change the name of your FASTA file from step 1 to the same name as the folder (except with .fasta at the end)
- Move the FASTA file into your new folder.
- Open terminal. Enter ‘cd *[Insert path to your new folder here]*’ to enter the correct directory

- You can drag the folder into the terminal window and it will enter the path to it automatically.
- Type in ‘makeblastdb -dbtype prot -in *[insert FASTA file name here]*’
- In the terminal it will tell you how many sequences were compiled into the database. Double check that this number is the **same as the number of search results from NCBI** when you downloaded the FASTA file.

- The download occasionally terminates early and doesn’t get every sequence, especially if you’re downloading multiple FASTA files at the same time.
- In the case that the numbers aren’t the same, try downloading the FASTA file again and repeat step 2.
- There should now be 3 new files in the folder, ending in .phr, .pin, and .psq
- You can remove the original FASTA file if you want to reduce storage consumption. Only the 3 newly generated files are necessary.
- You now have a properly formatted BLAST database.

### Step 3: Insert the new folder into the database file tree

- In the ProteoSync folder, there’s a folder called databases. In this folder is another folder called species, which contains all the species-specific databases.
- The directories in the species folder are organized in a tree structure by taxonomy. The GUI uses this file structure to generate the taxonomy settings window.
- Insert your new folder into the file tree where it belongs taxonomically. Your species should then appear on the taxonomy menu the next time you open the application.
- NOTE: You can expand the taxonomic tree structure to include more taxonomic groups by creating new directories to organize your species folders. These groups will also appear on the taxonomy settings menu.

- Make sure folder names don’t contain spaces or unnecessary special characters
- Make sure that individual species database folders contain **no other directories**. If a directory contains any subdirectories, it is assumed to be a taxonomic group folder, and the species database will not be found.

## Acknowledgments

The authors wish to acknowledge members of the Sicheri lab for help with program testing.

## Author contributions

E.S., D.M. and F.S conceived the project and wrote the manuscript. E.S. wrote the program. F.S. secured funding and supervised the project.

## Statements and declarations

### Ethical considerations

Not applicable

### Consent to participate

Not applicable

### Consent for publication

Not applicable

### Data availability

The Proteosync program, as well as a set of starter species databases, is available for download on the Sicheri lab website and on GitHub

## Figure Legends

**Supplementary Information Figure 1.**
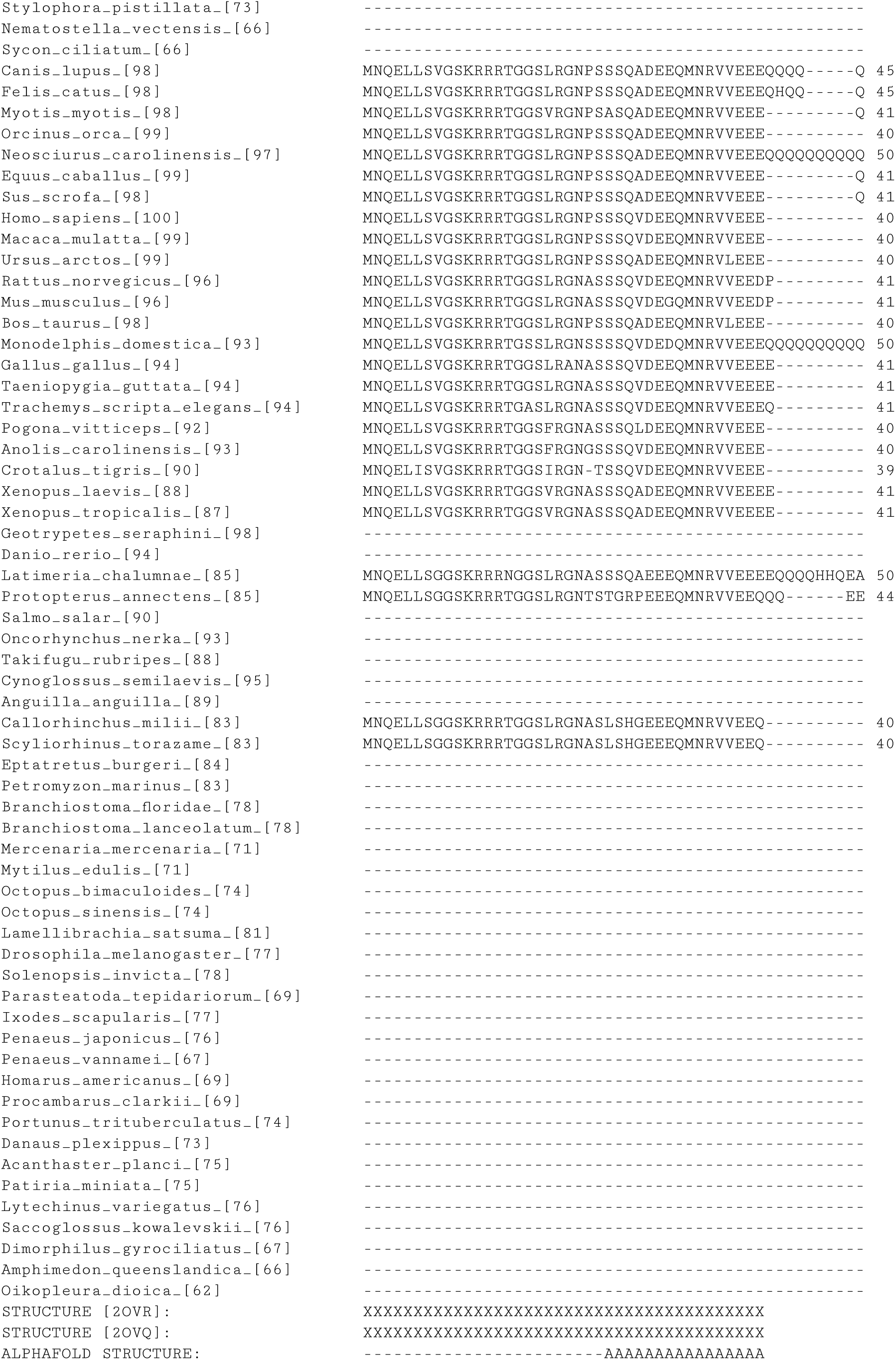
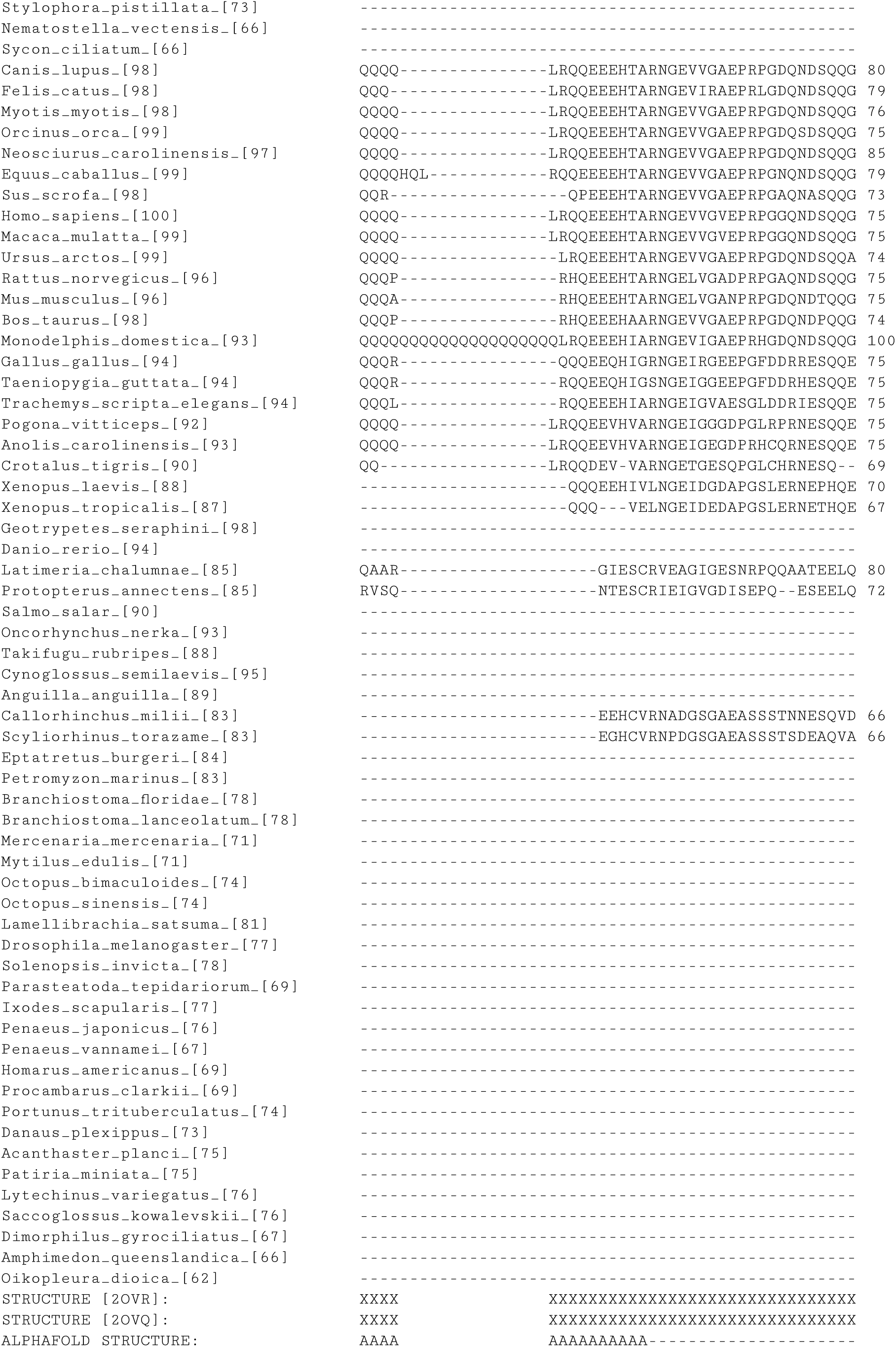
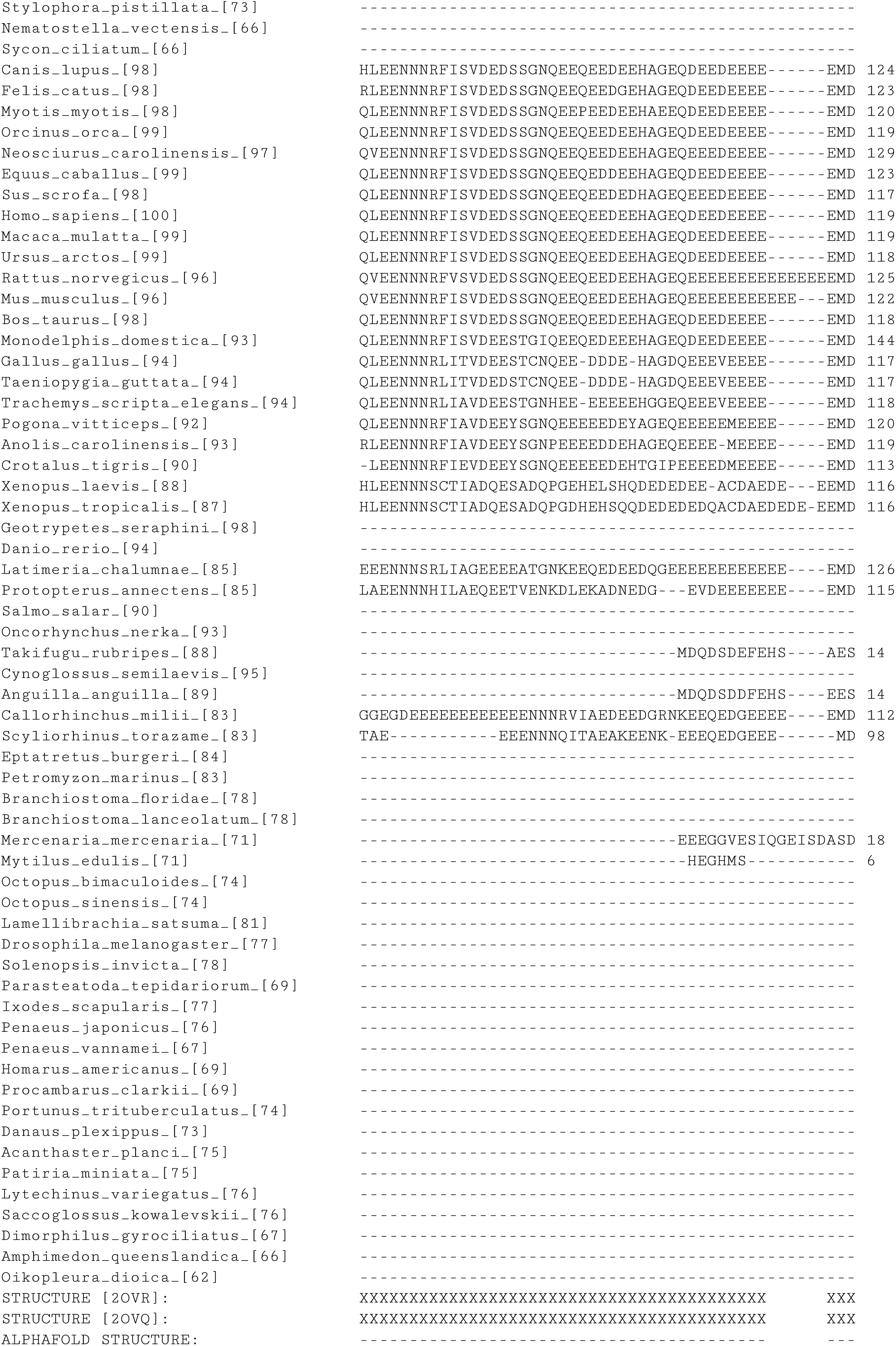
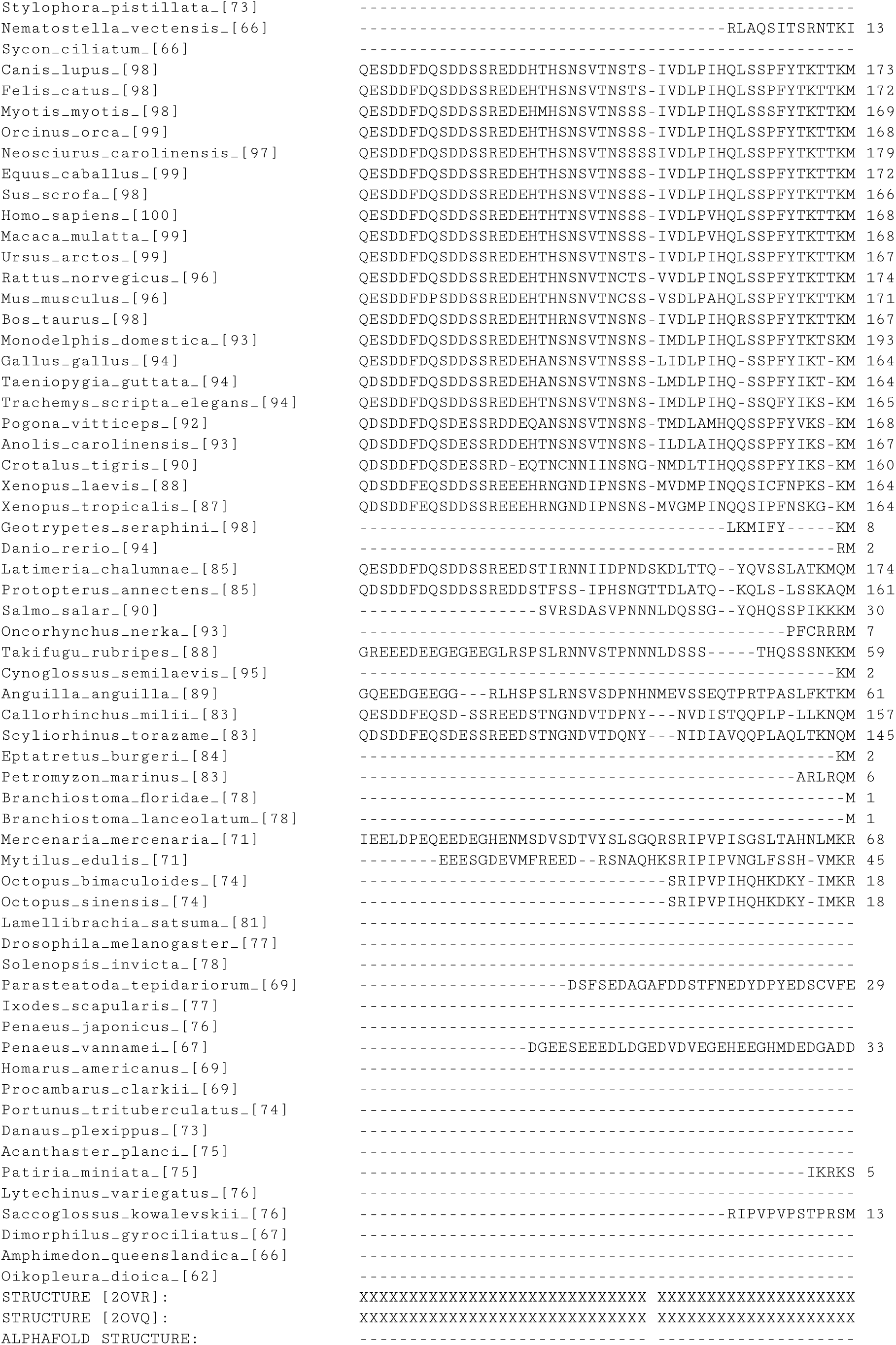
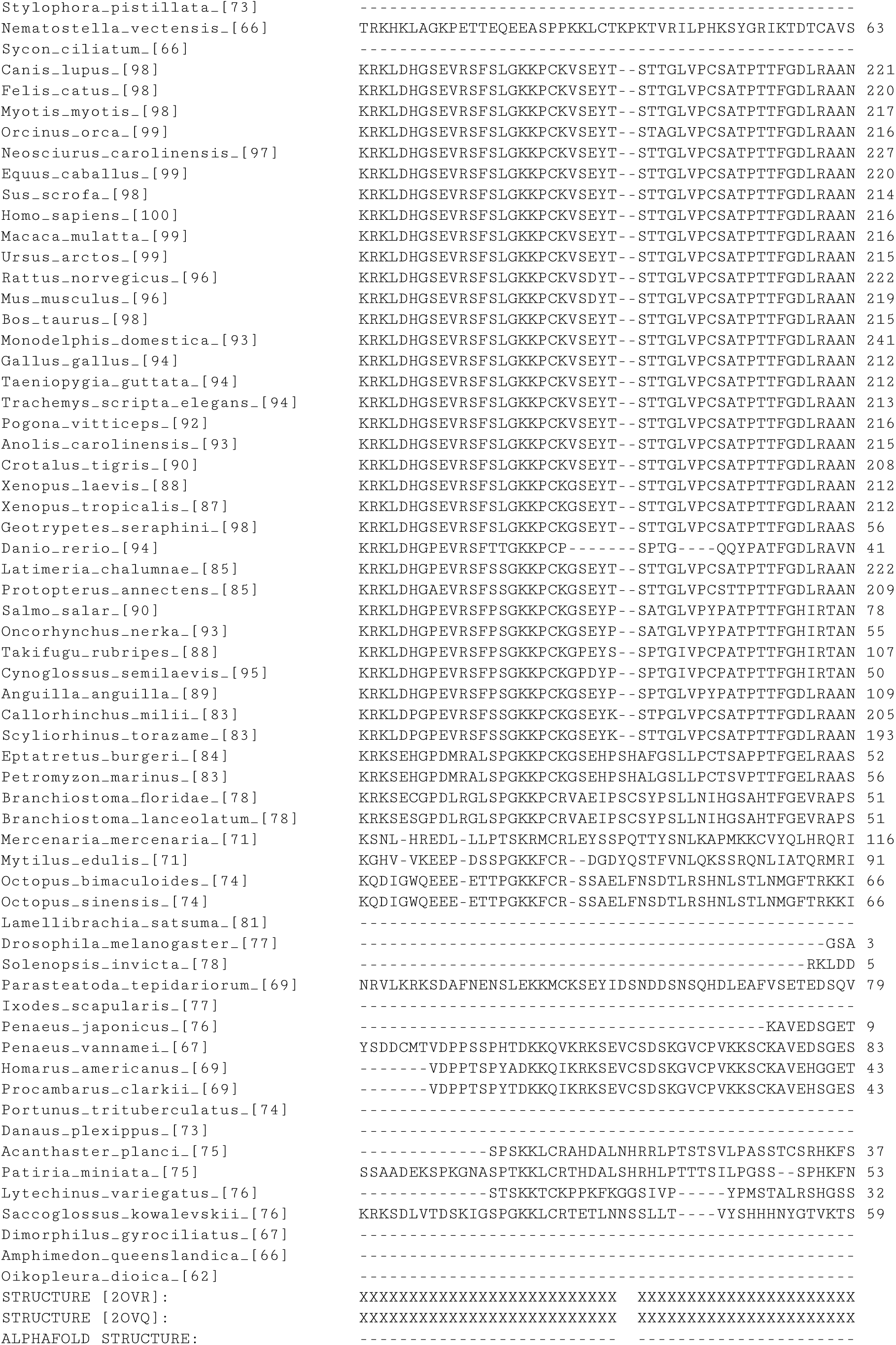
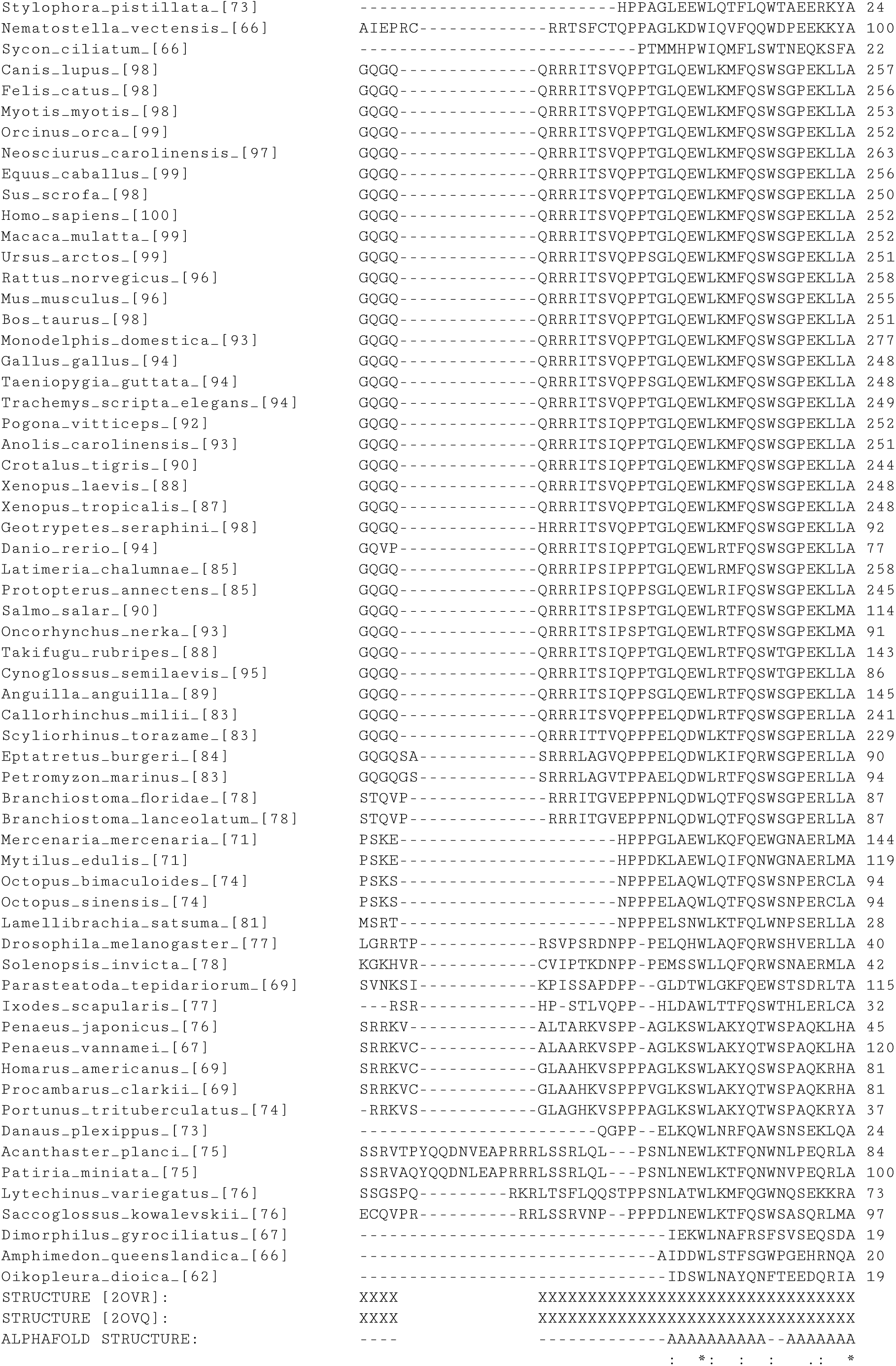
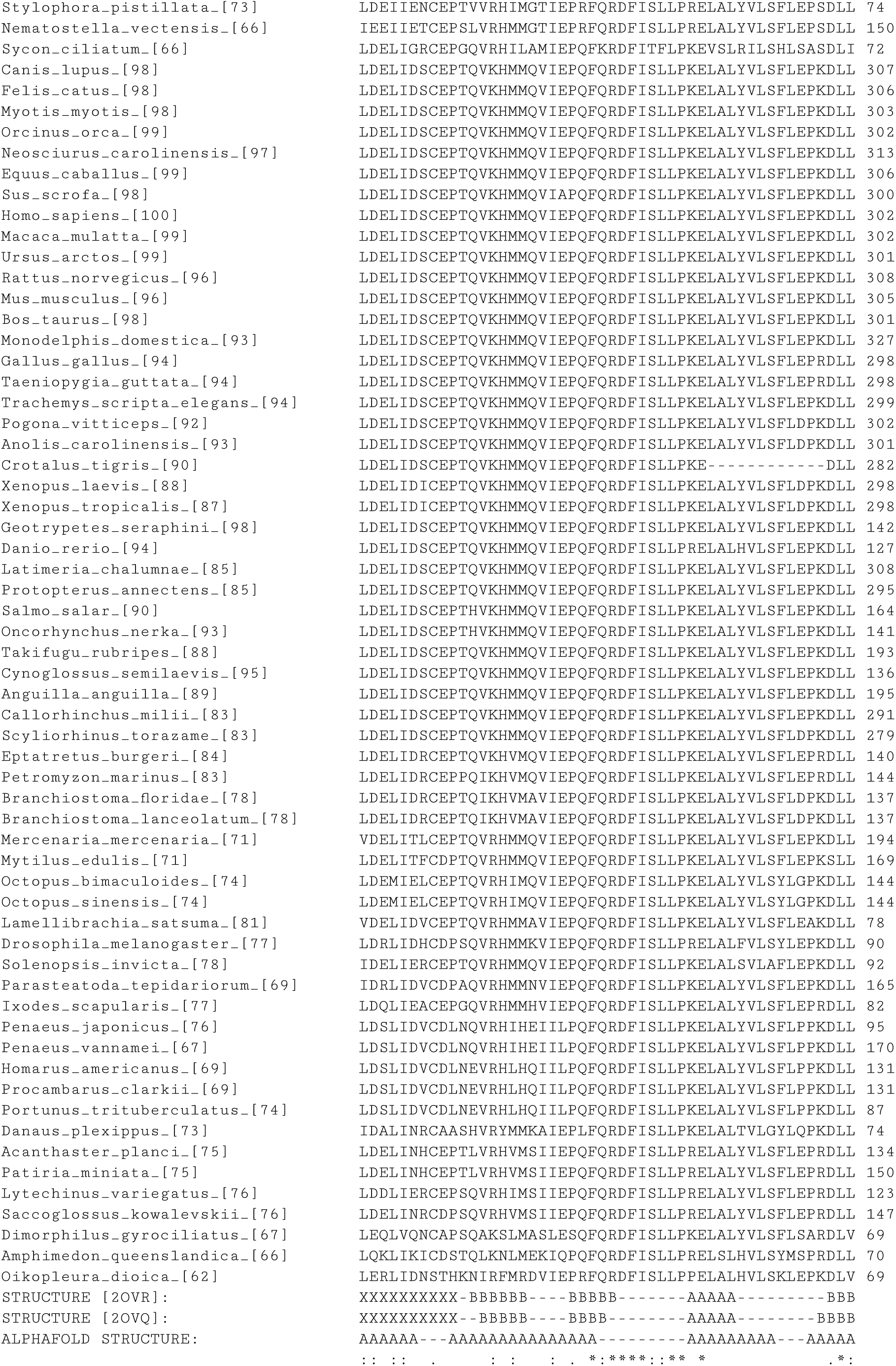
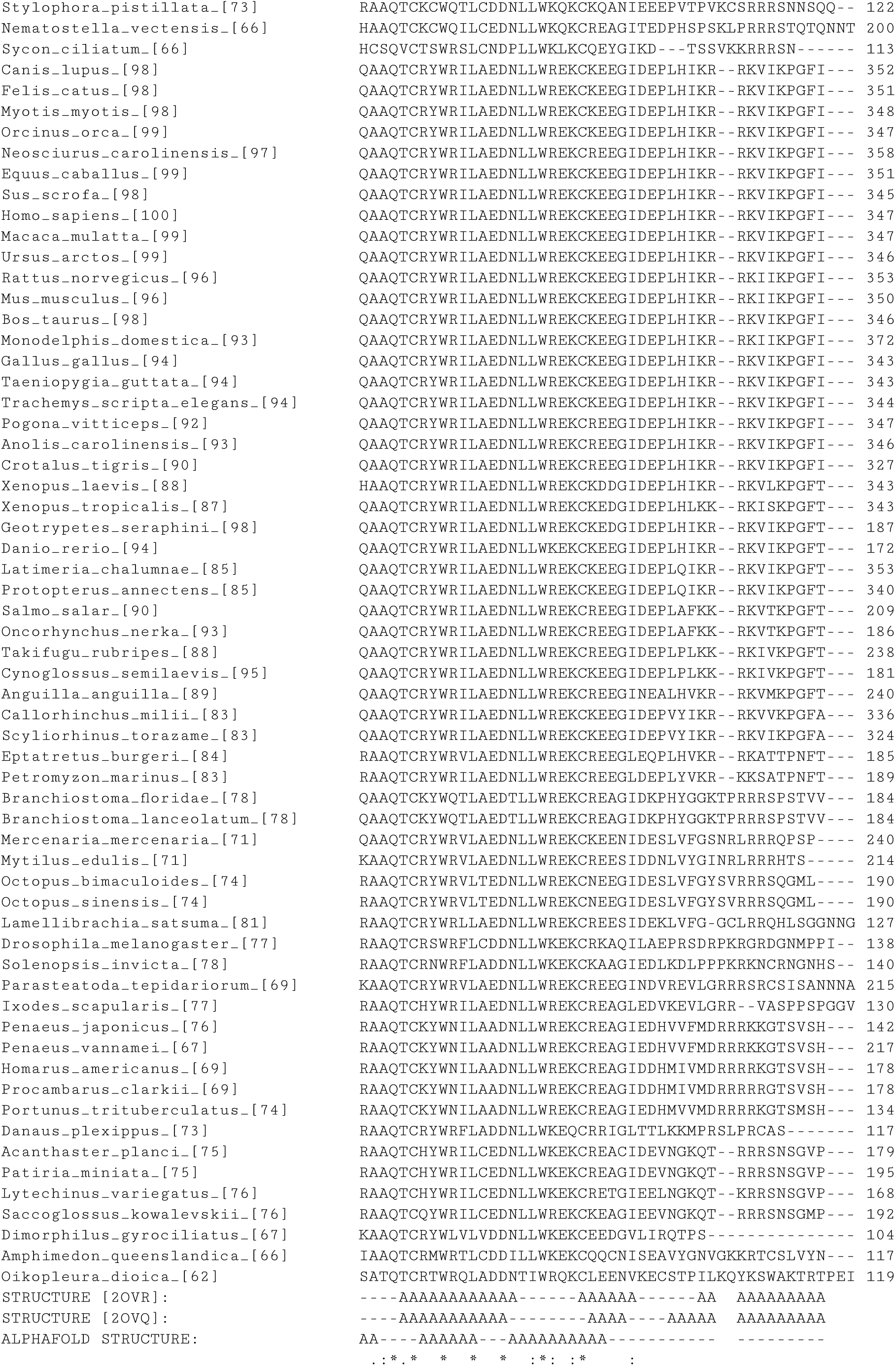
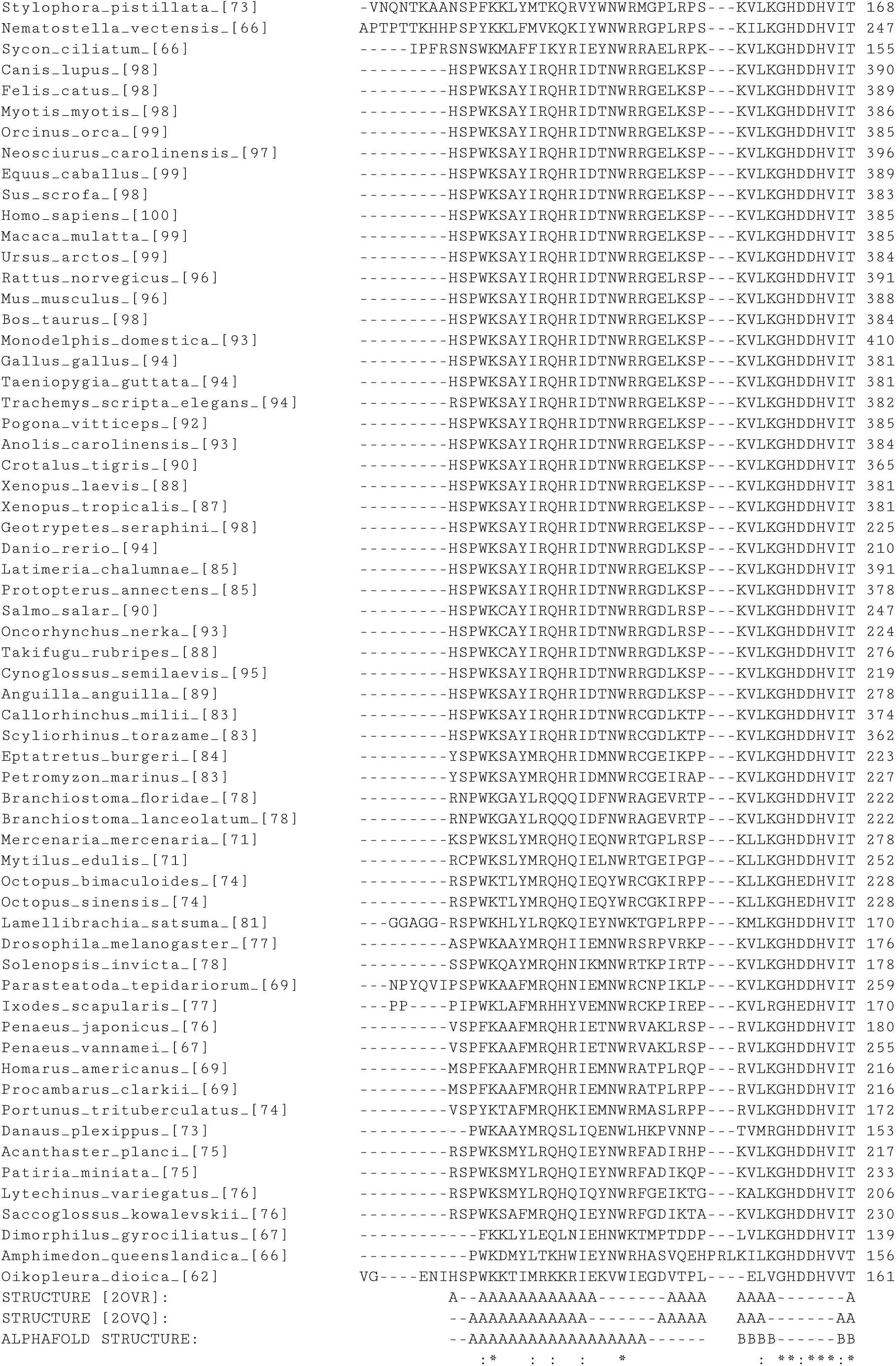
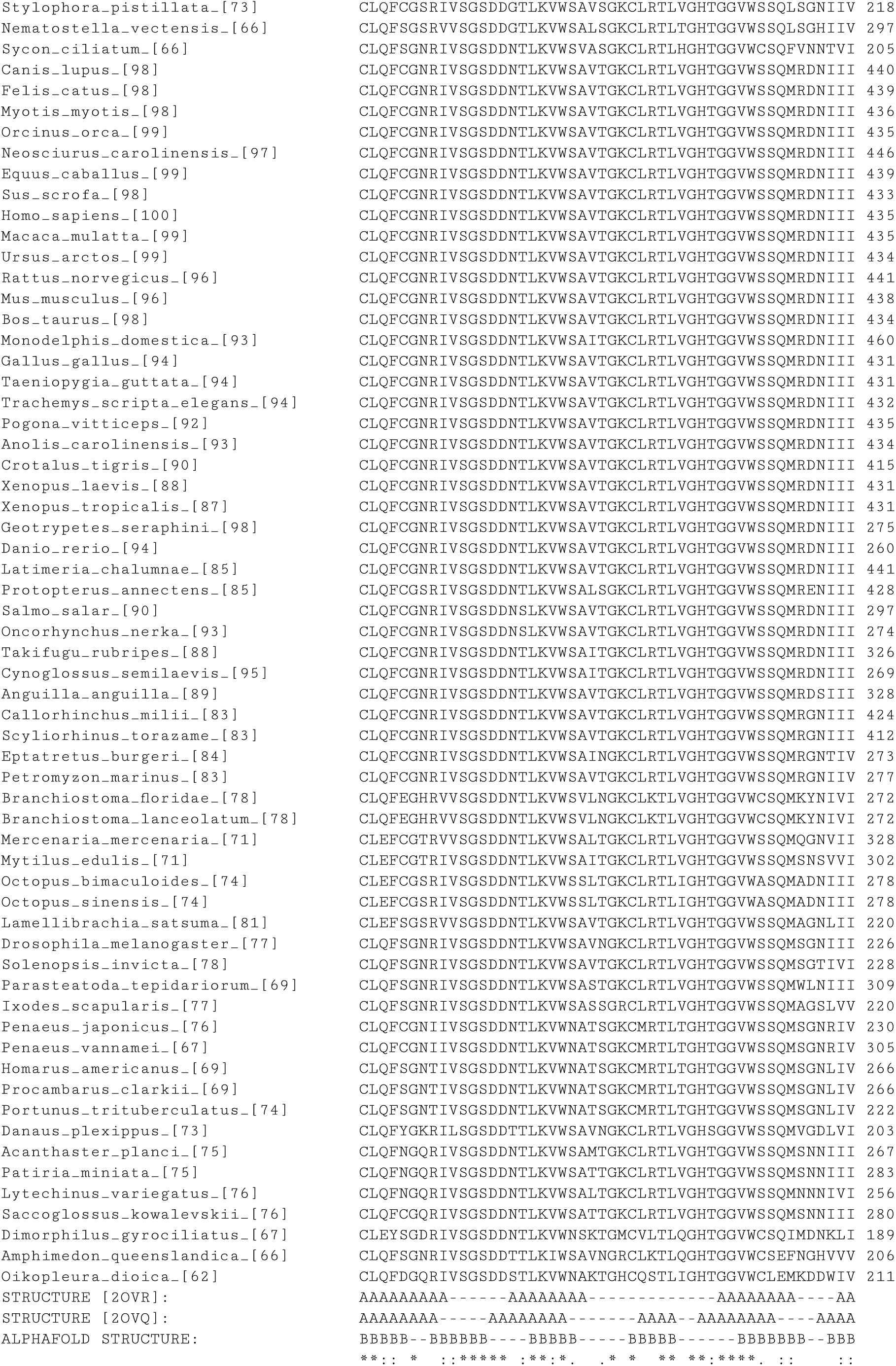
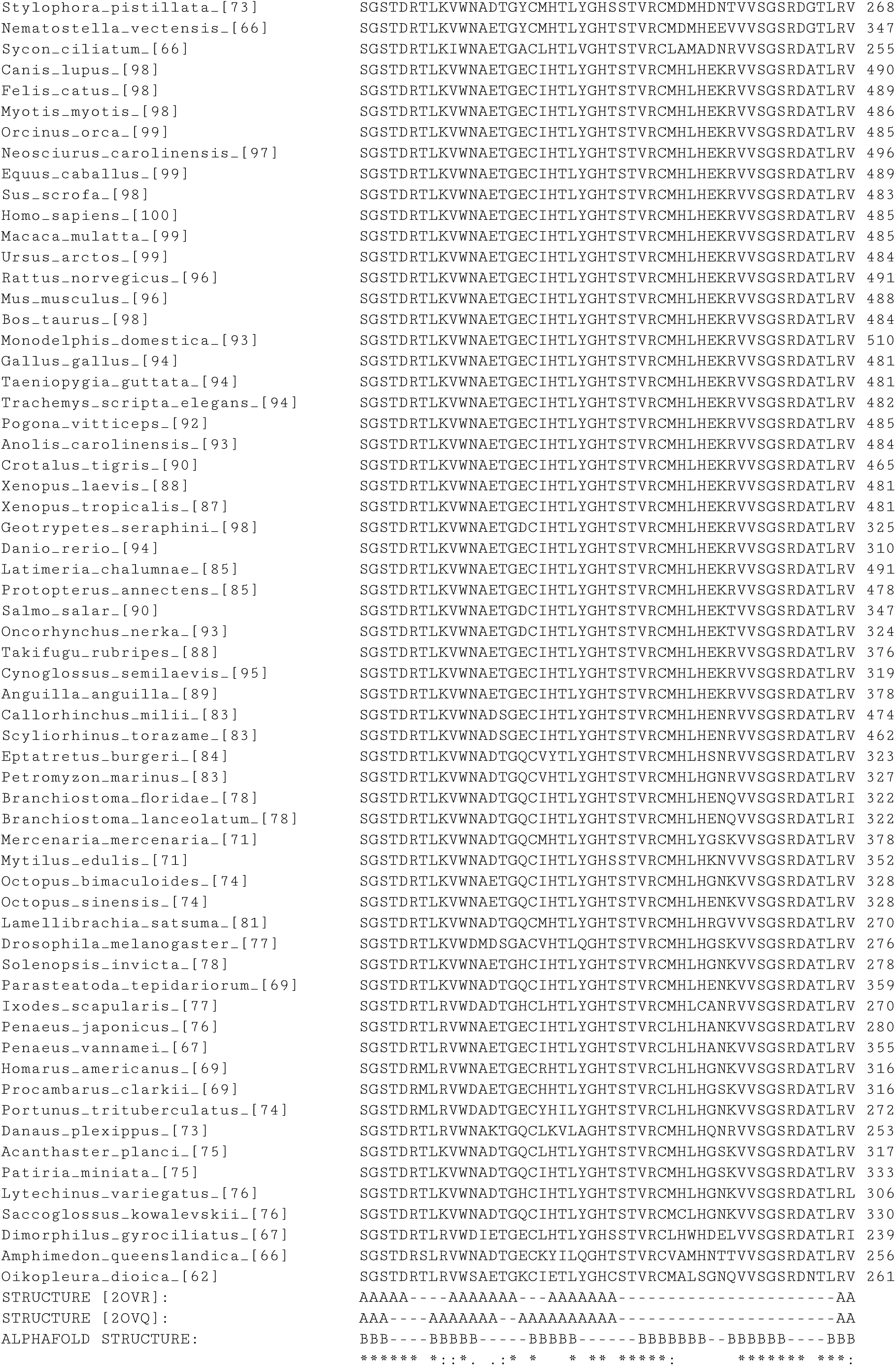
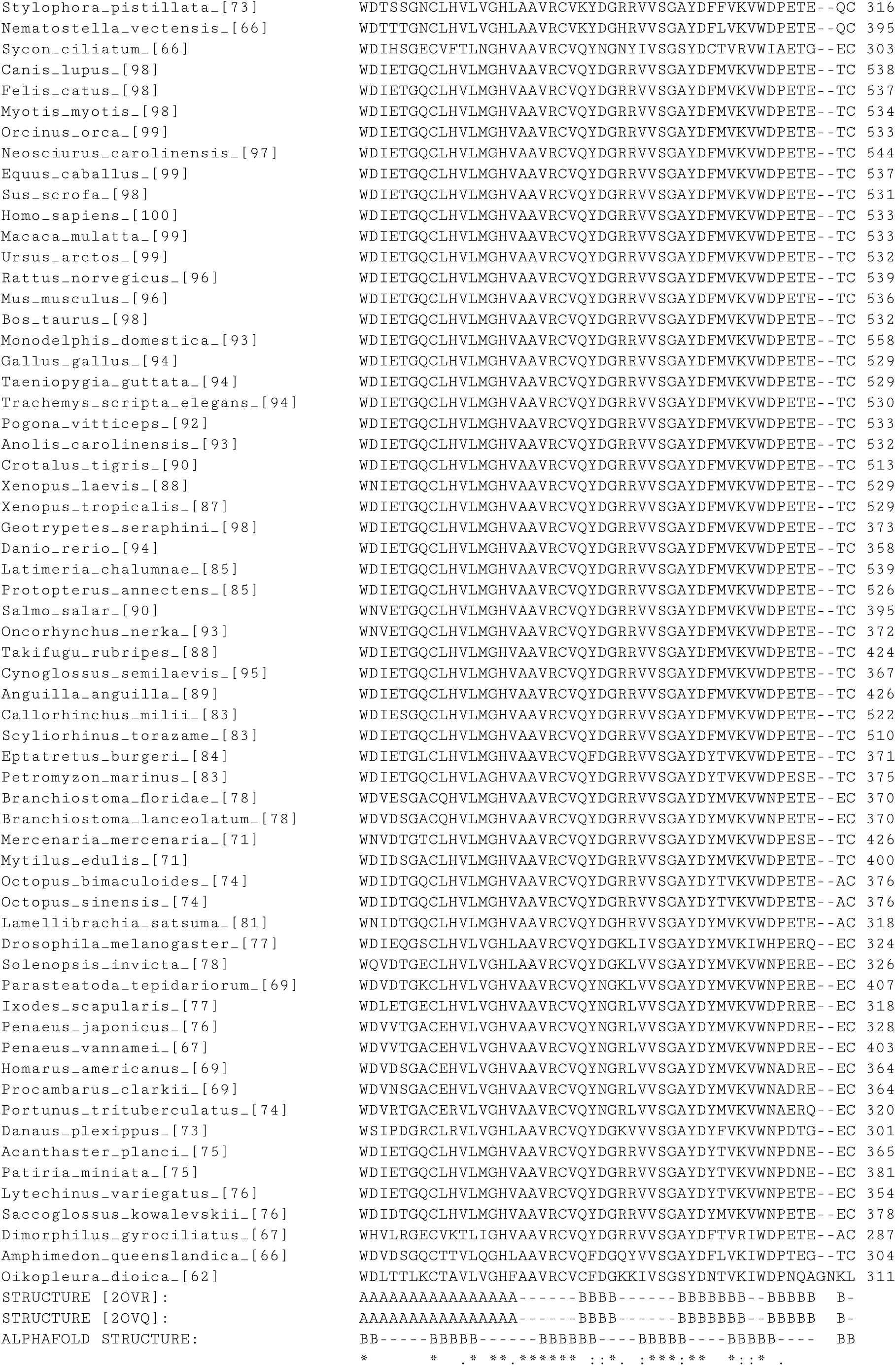
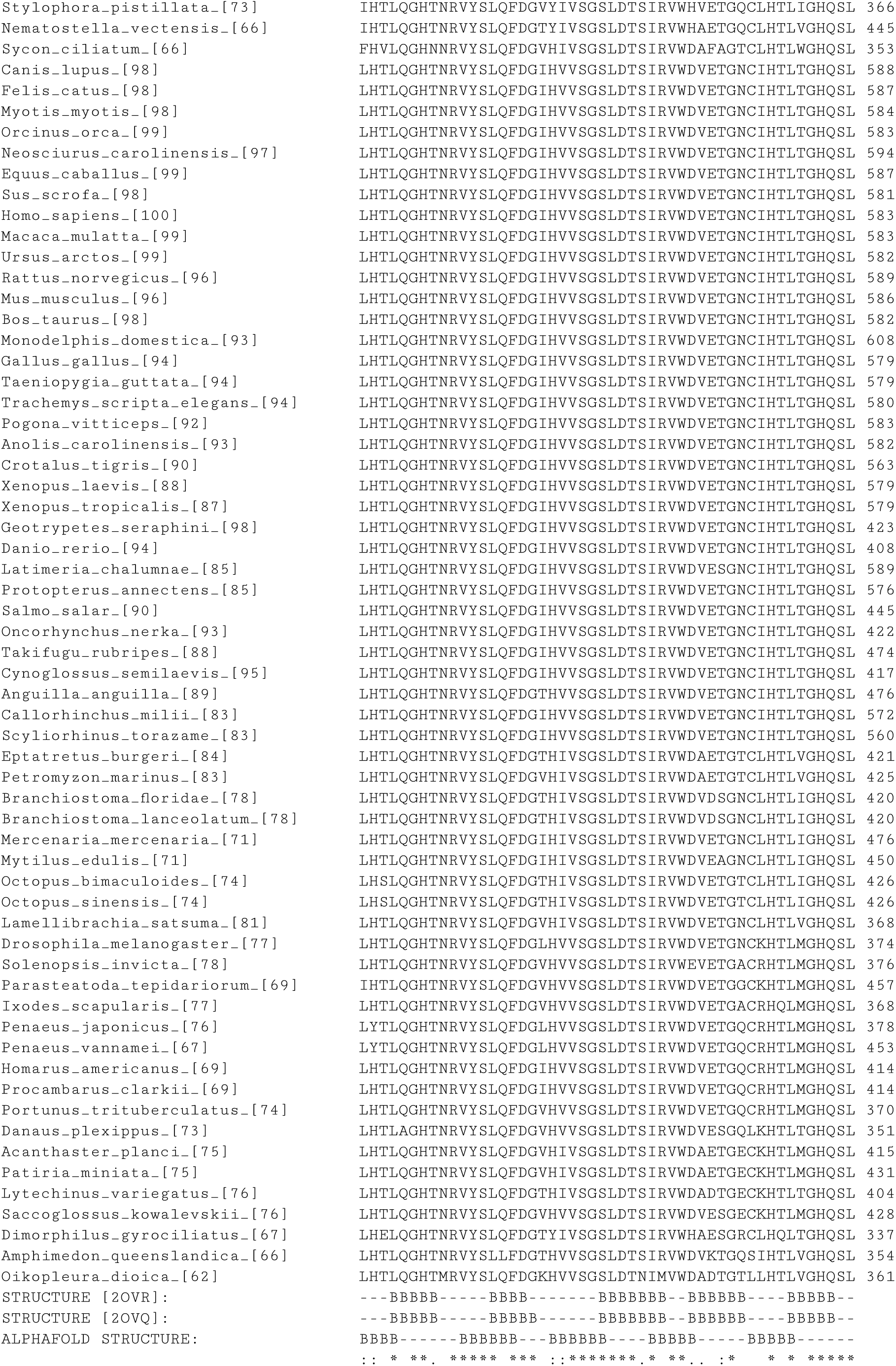
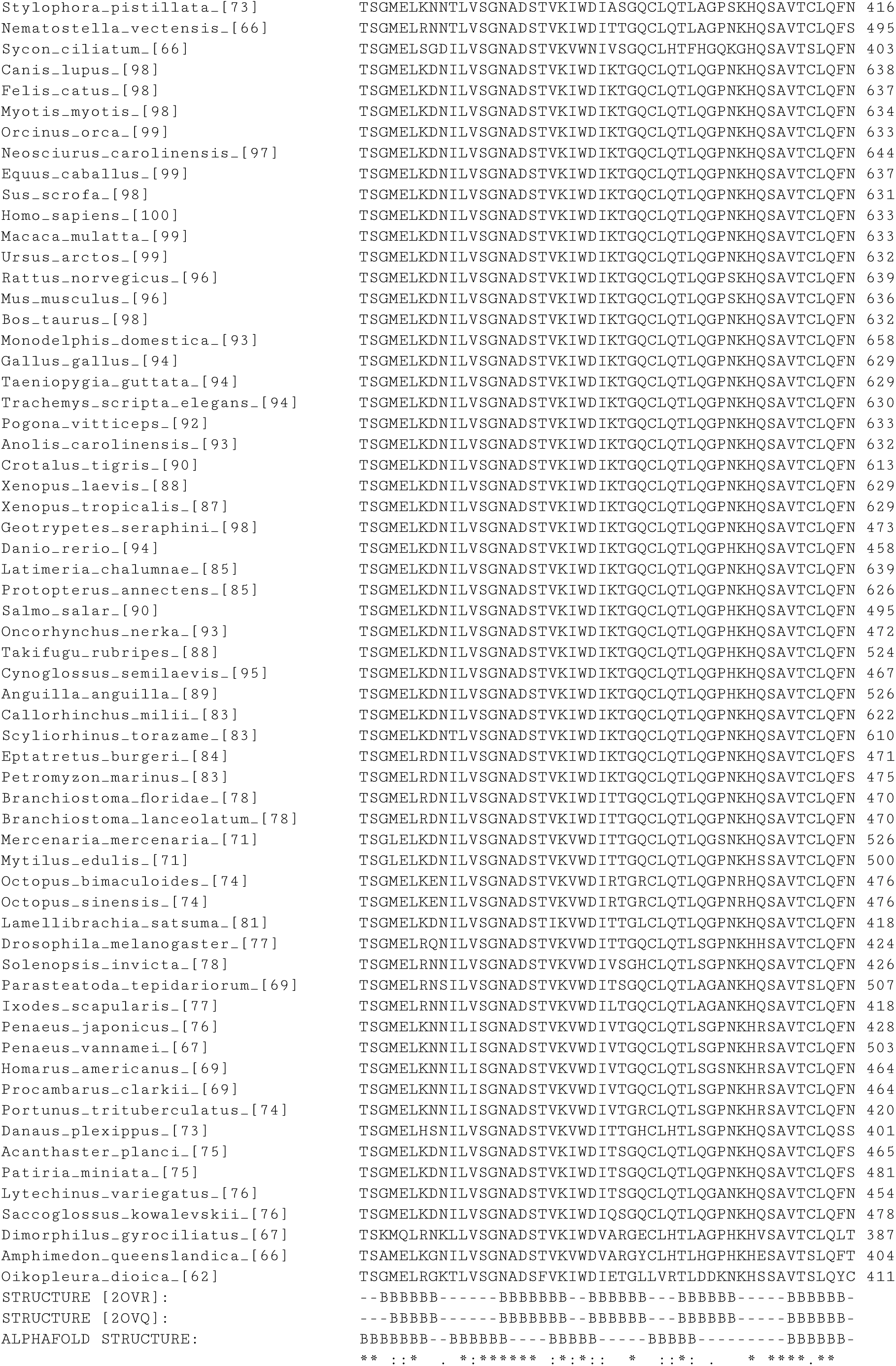
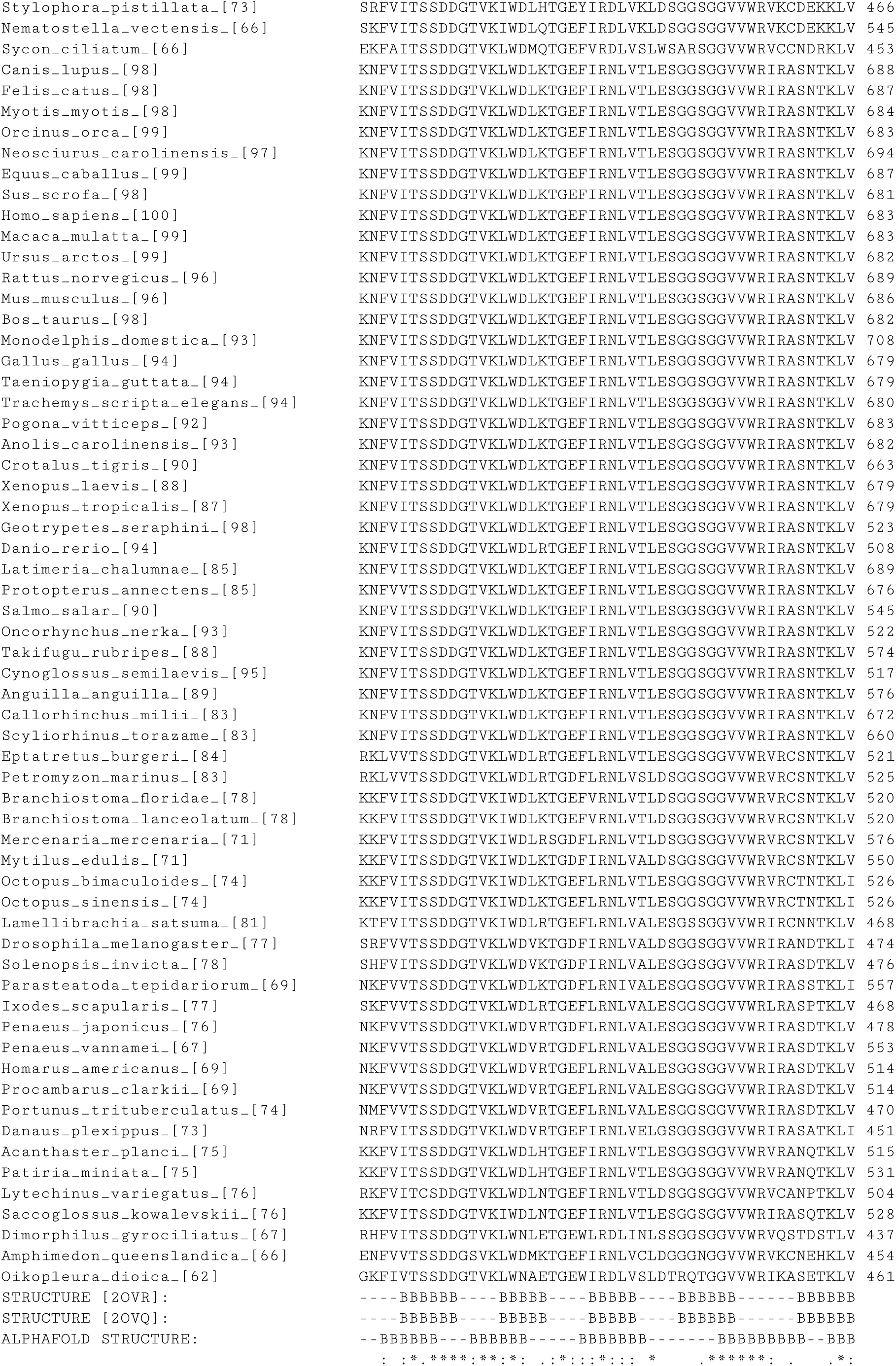
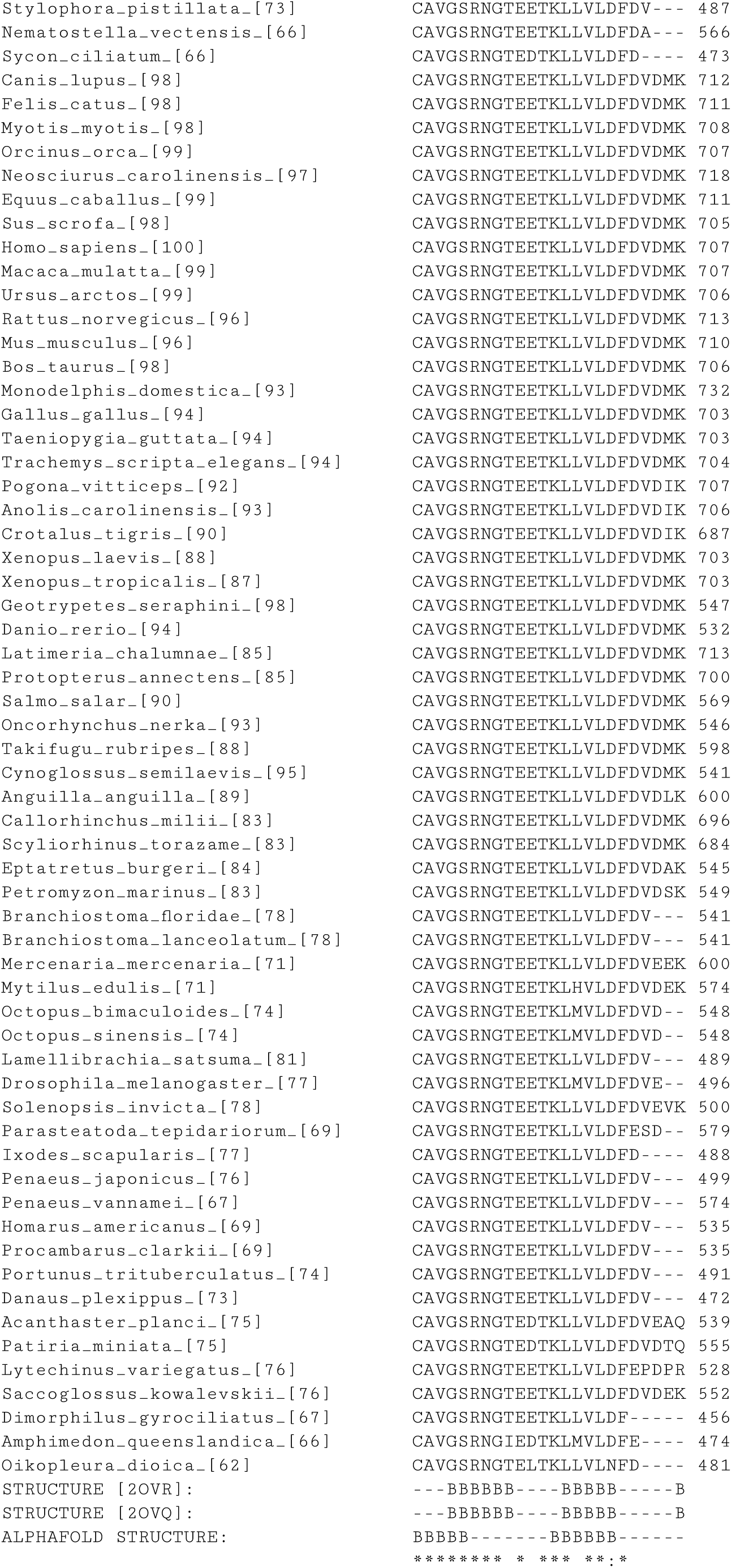
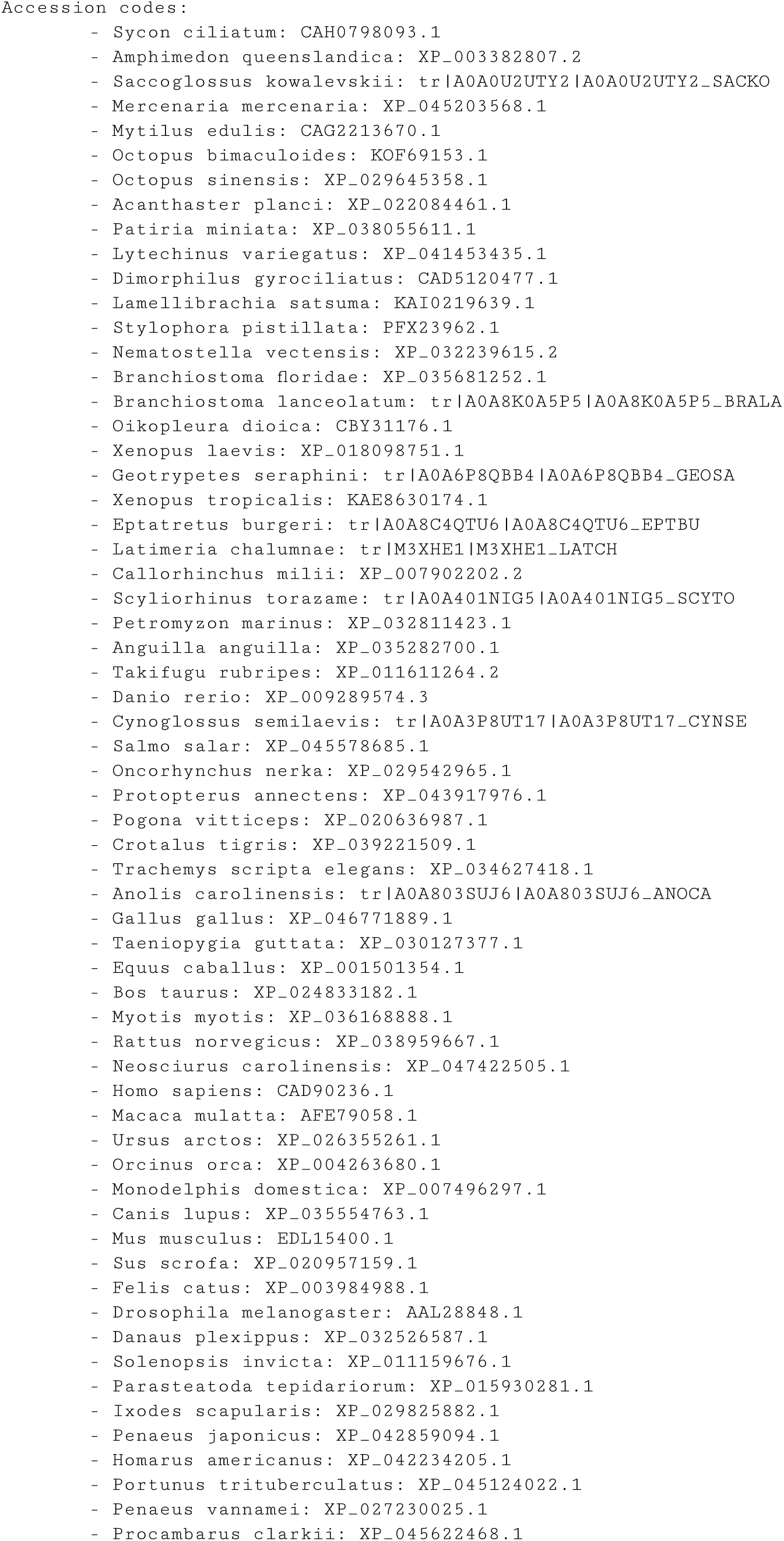
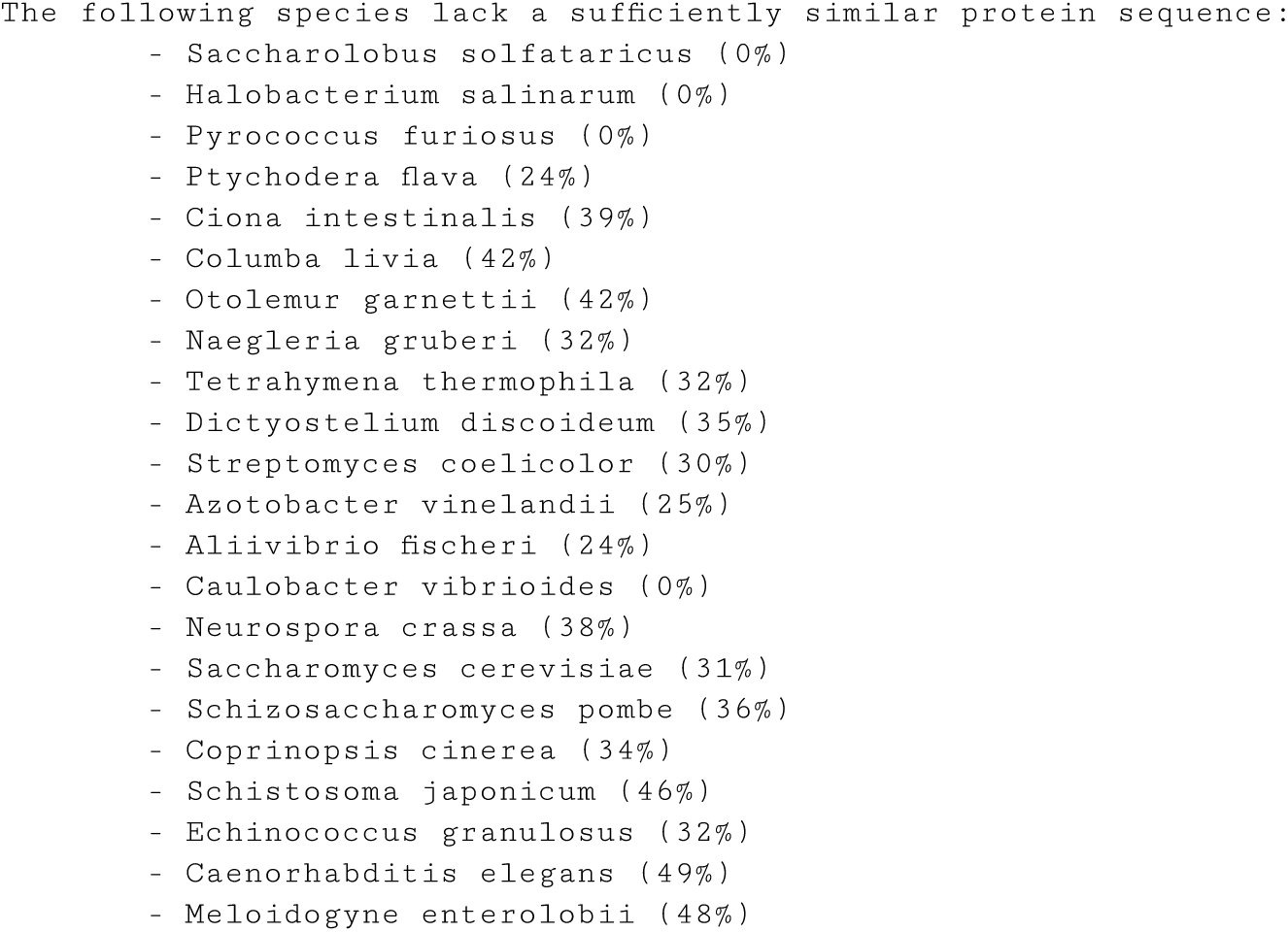
ProteoSync sequence alignment output file from the analysis of human FBXW7.

**Supplementary Information Figure 2.**
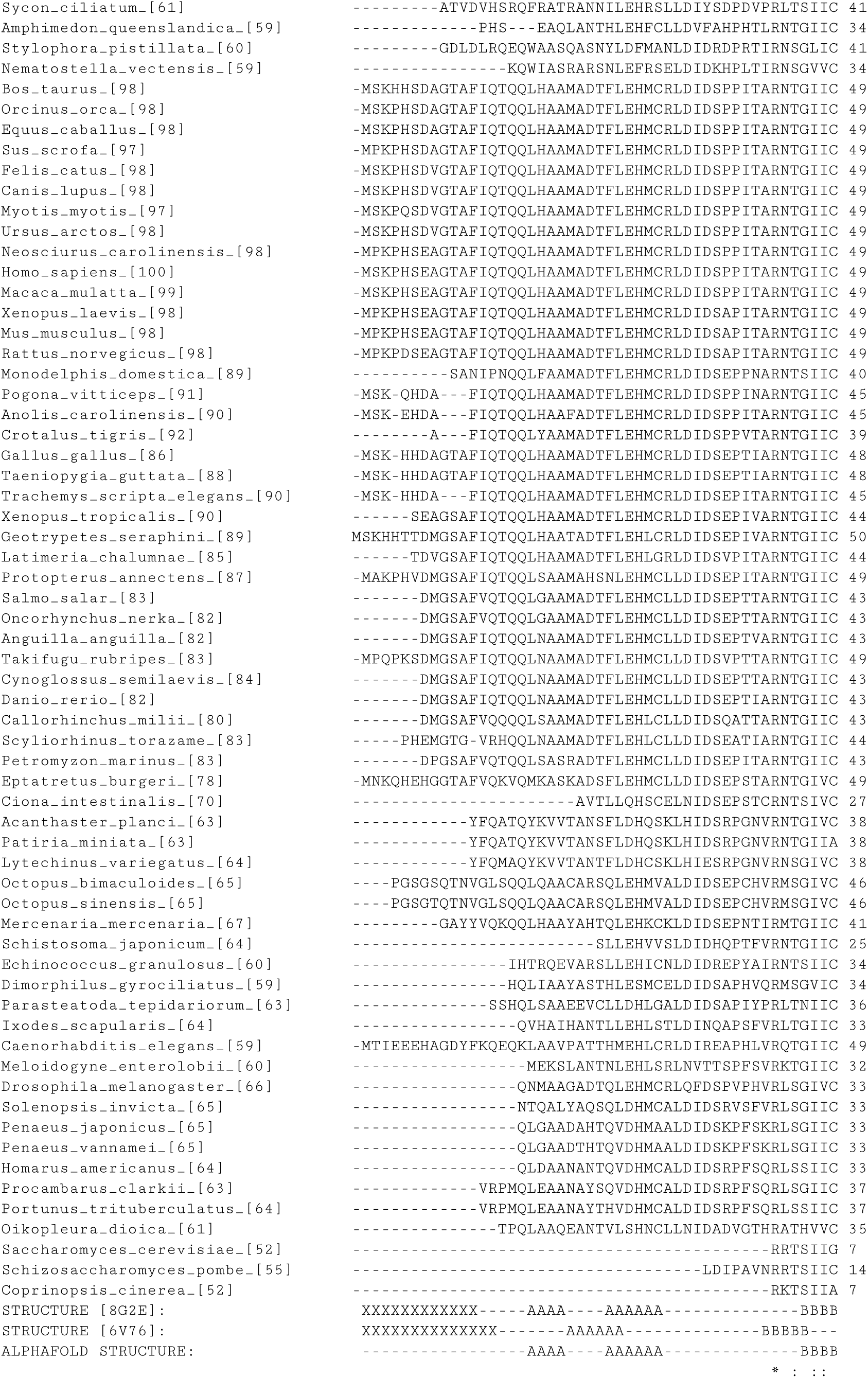
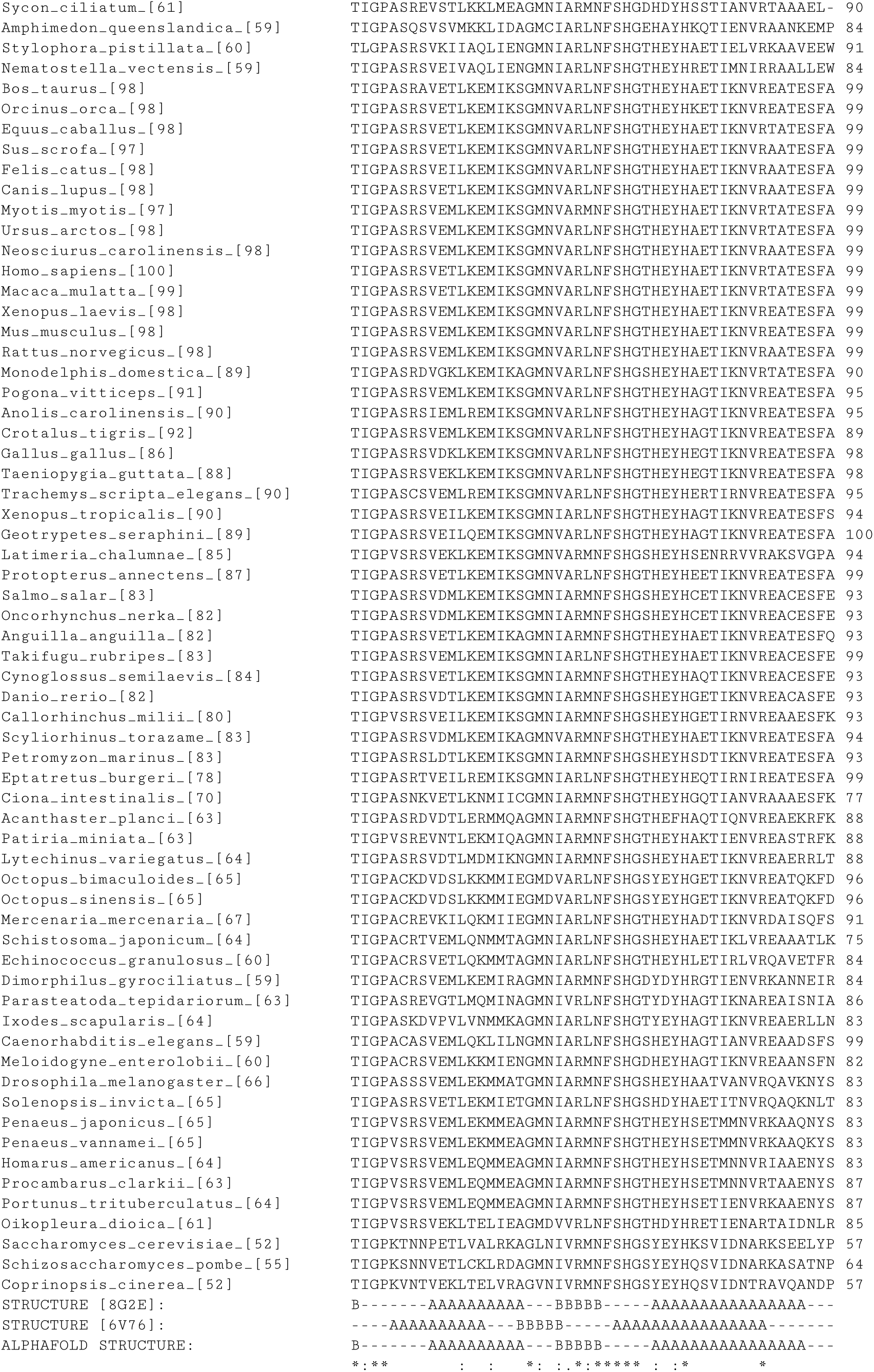
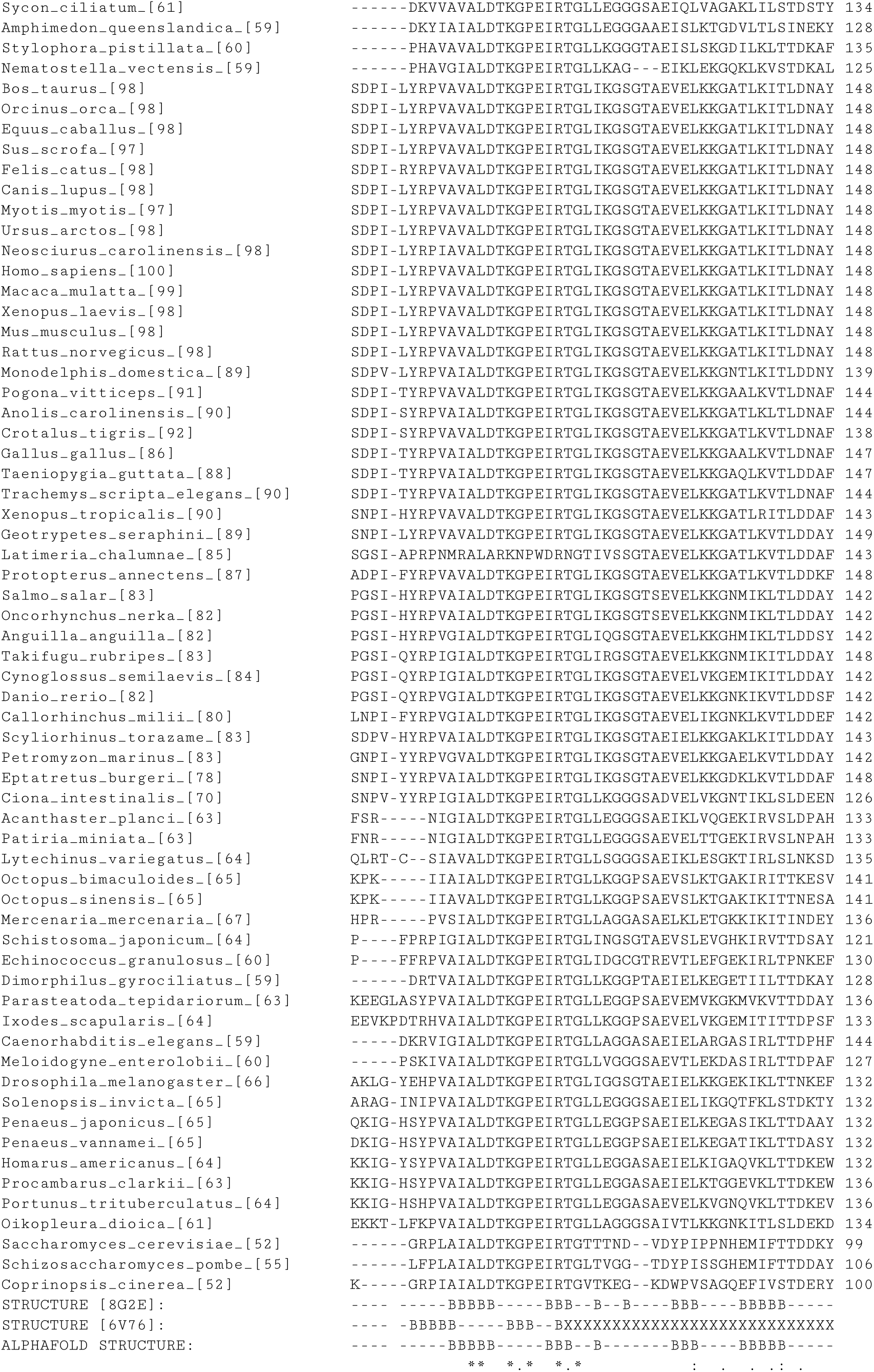
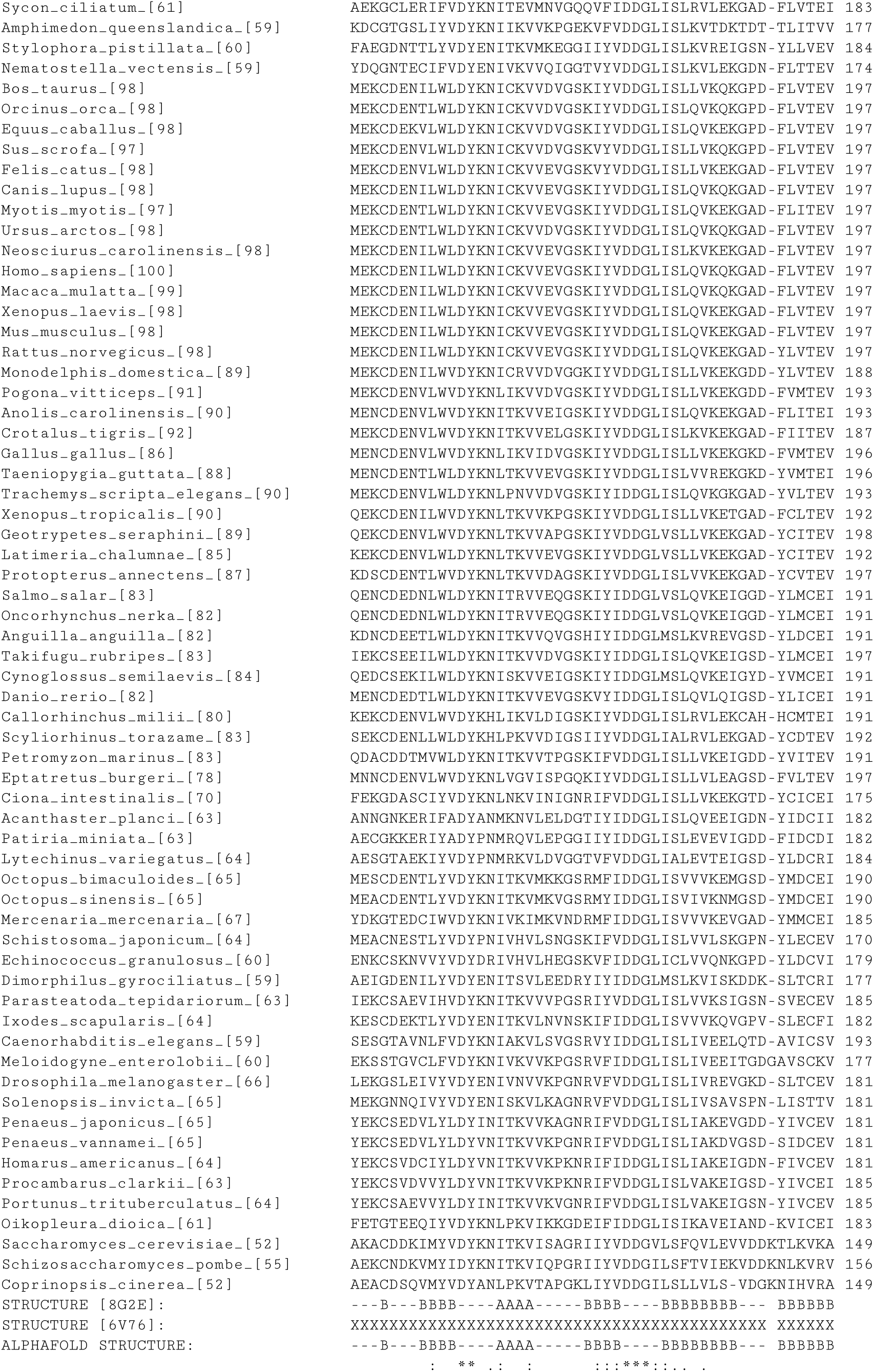
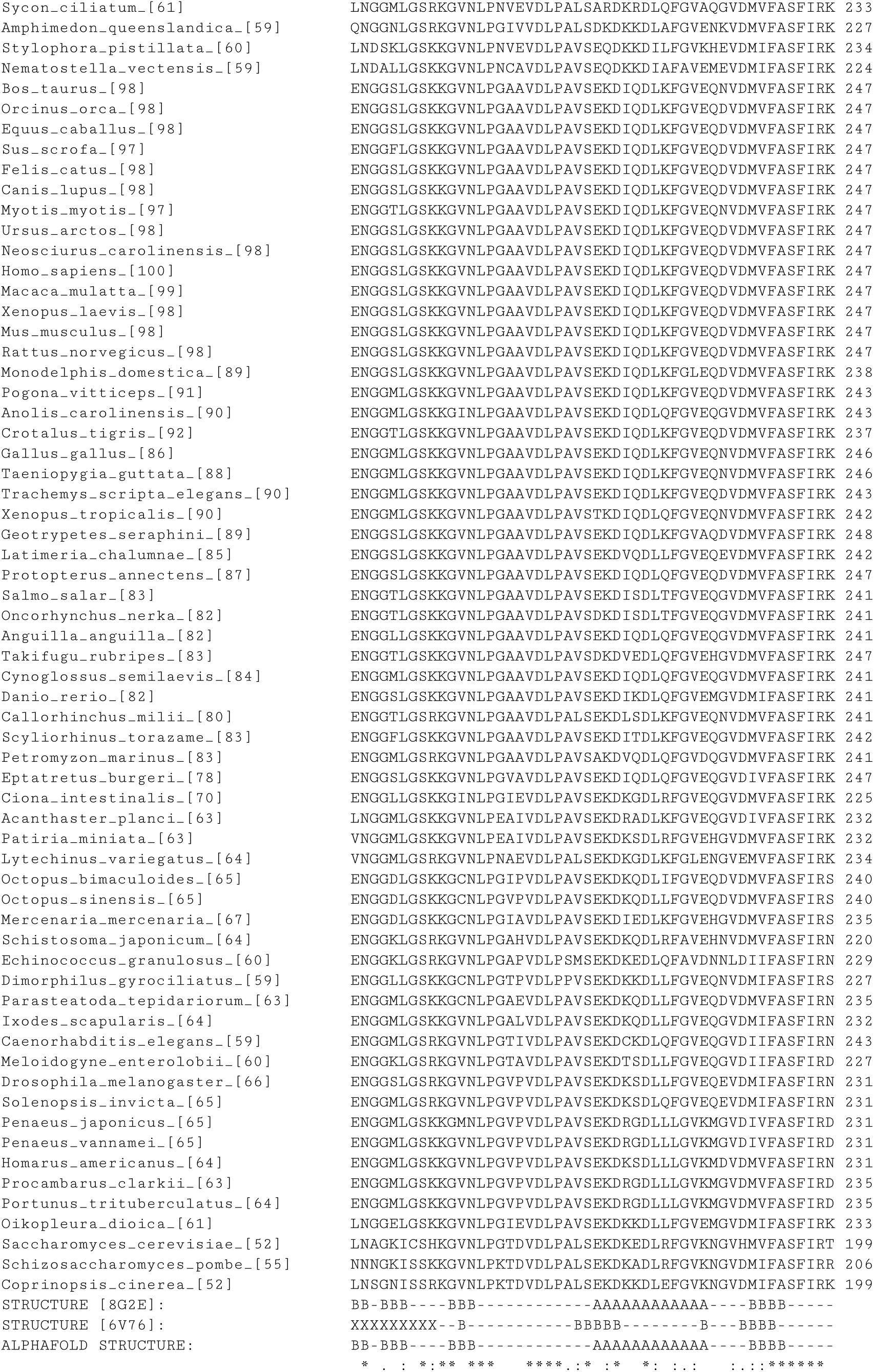
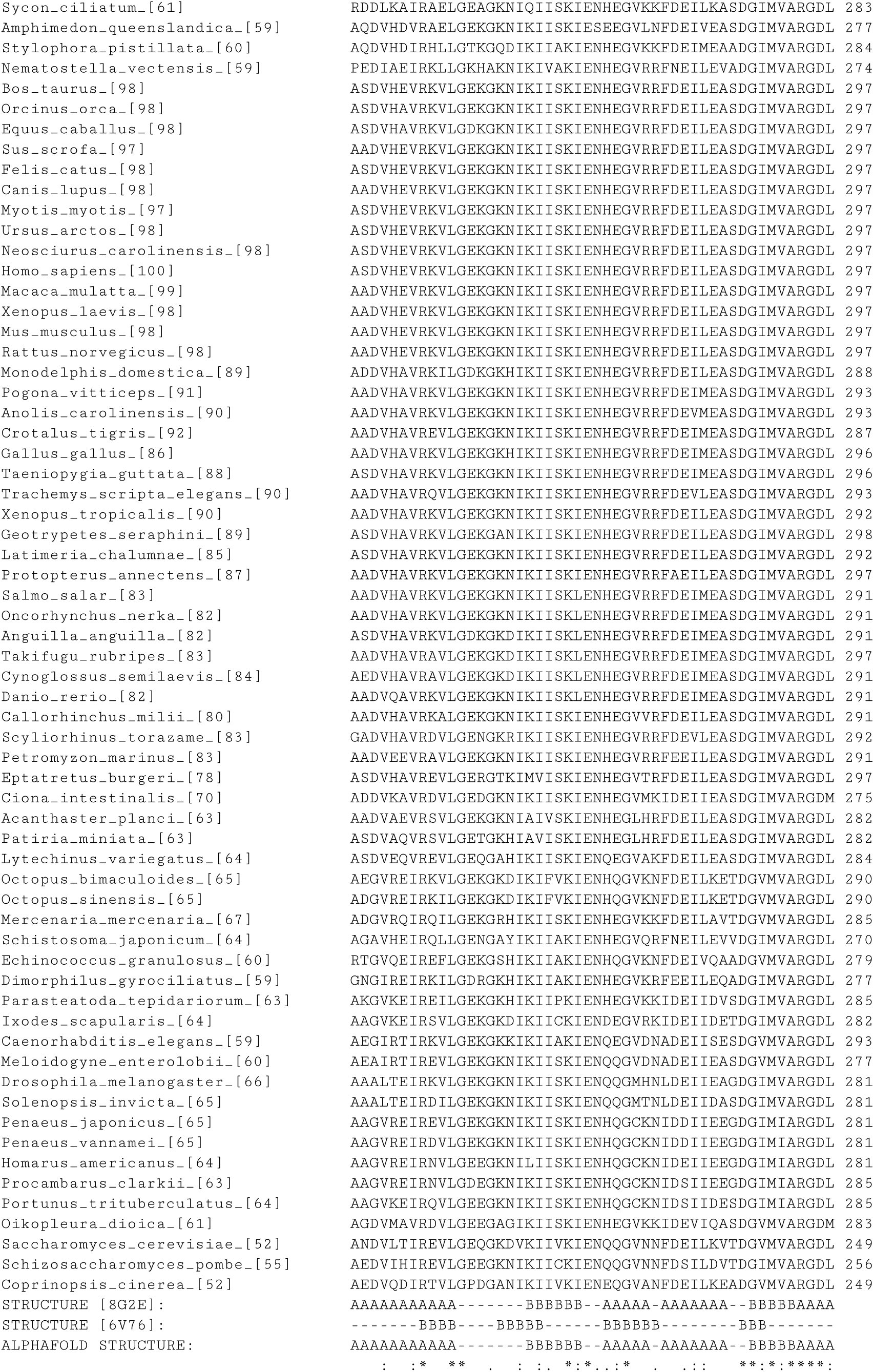
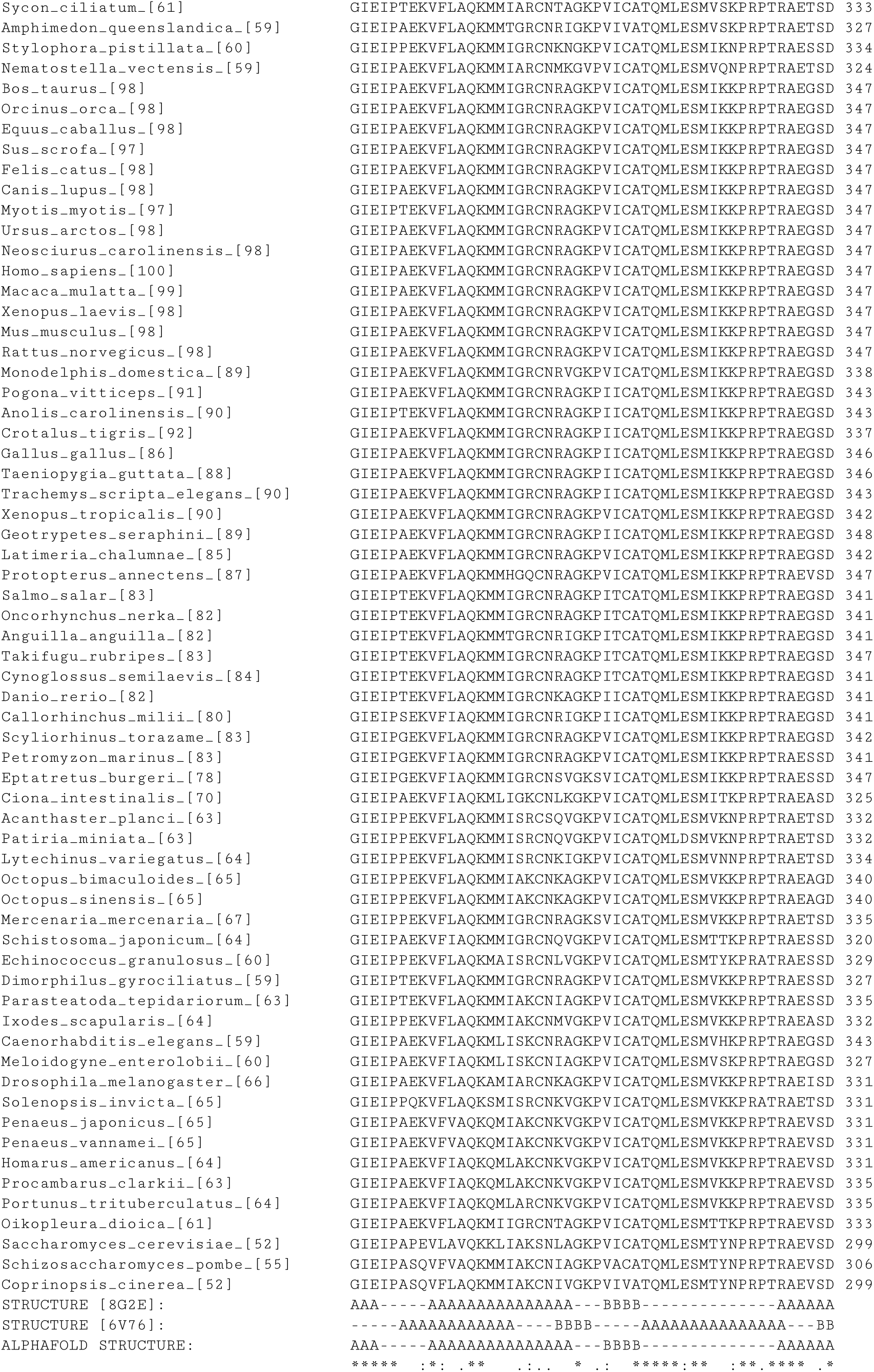
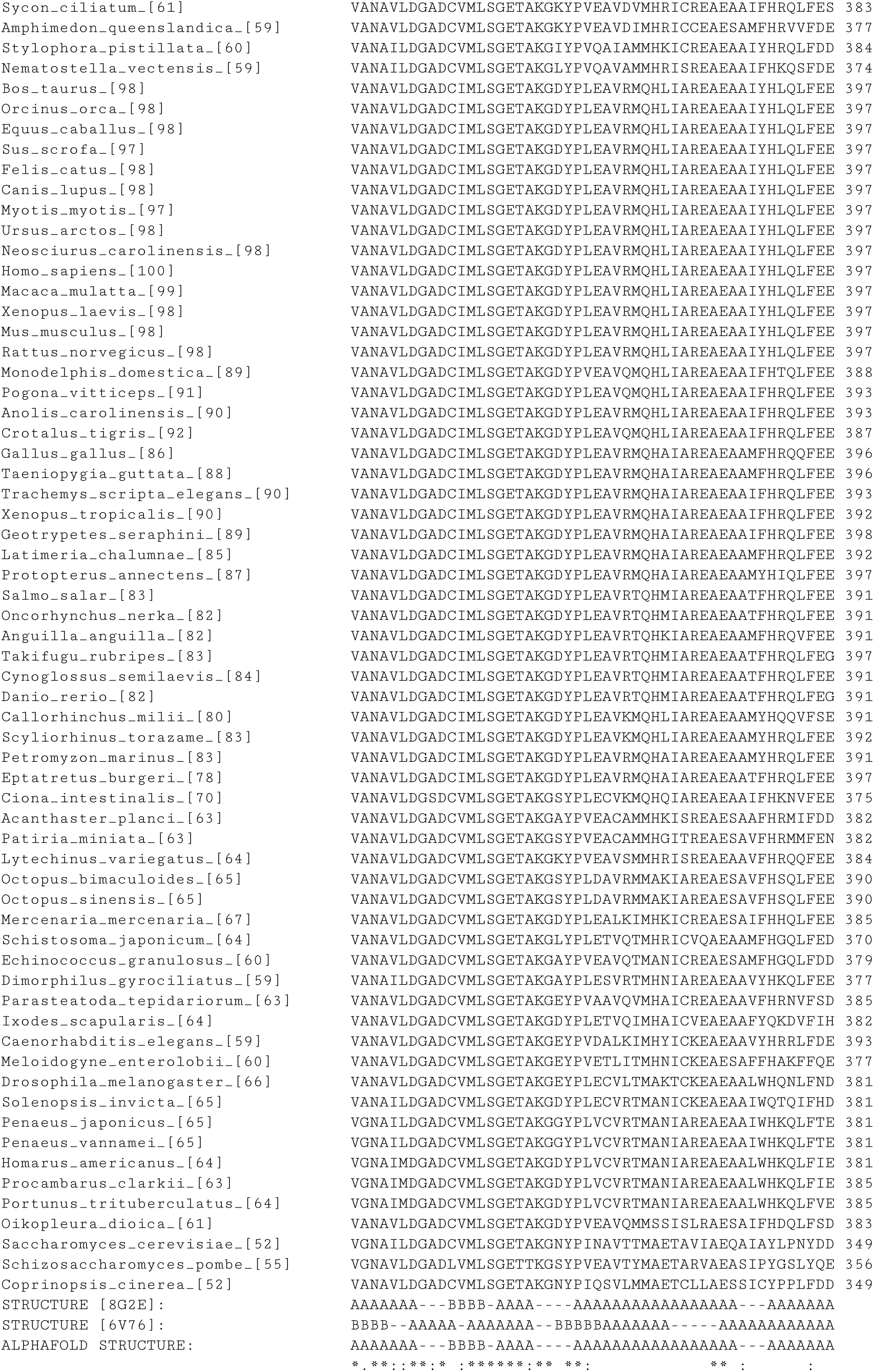
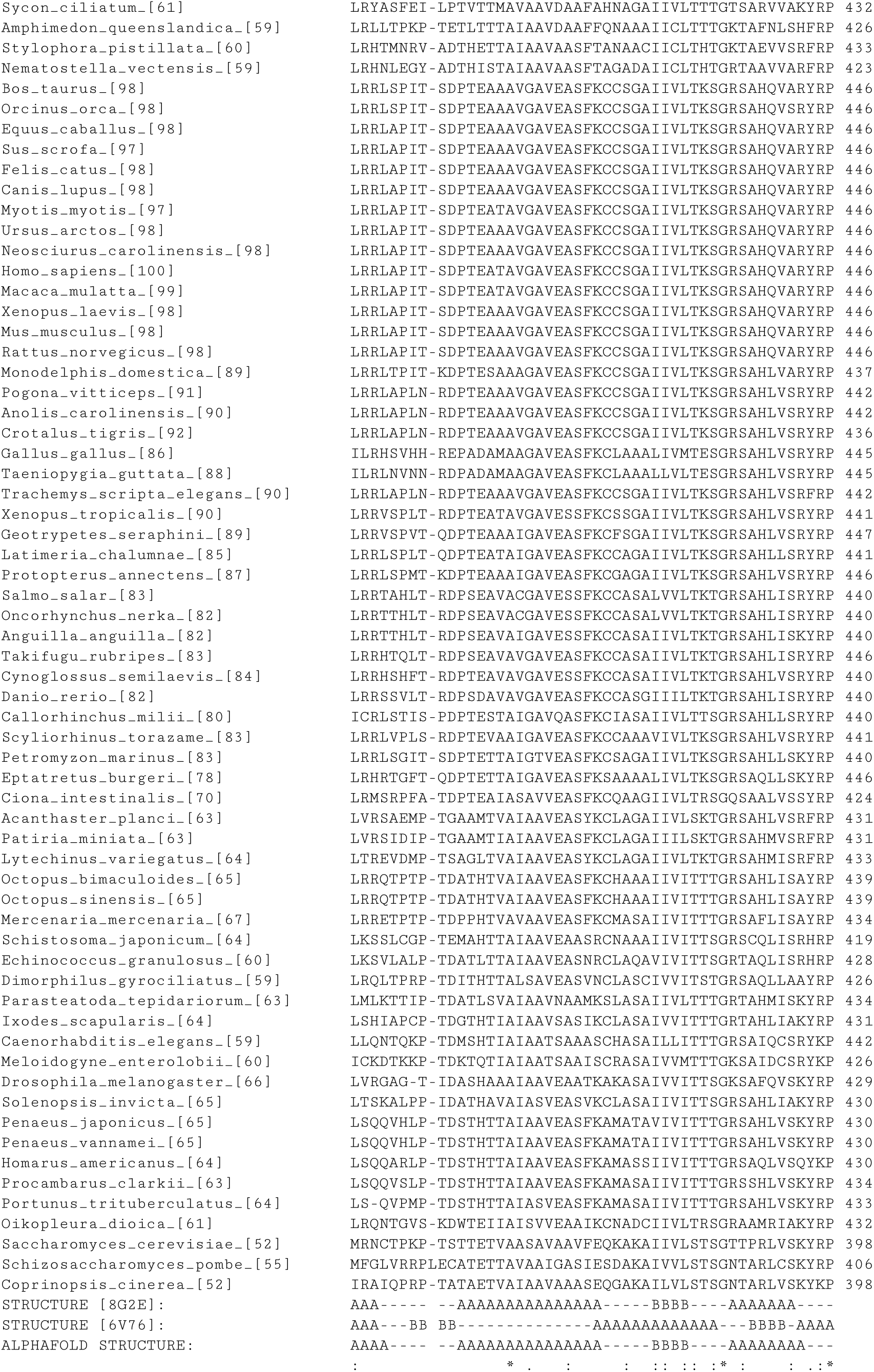
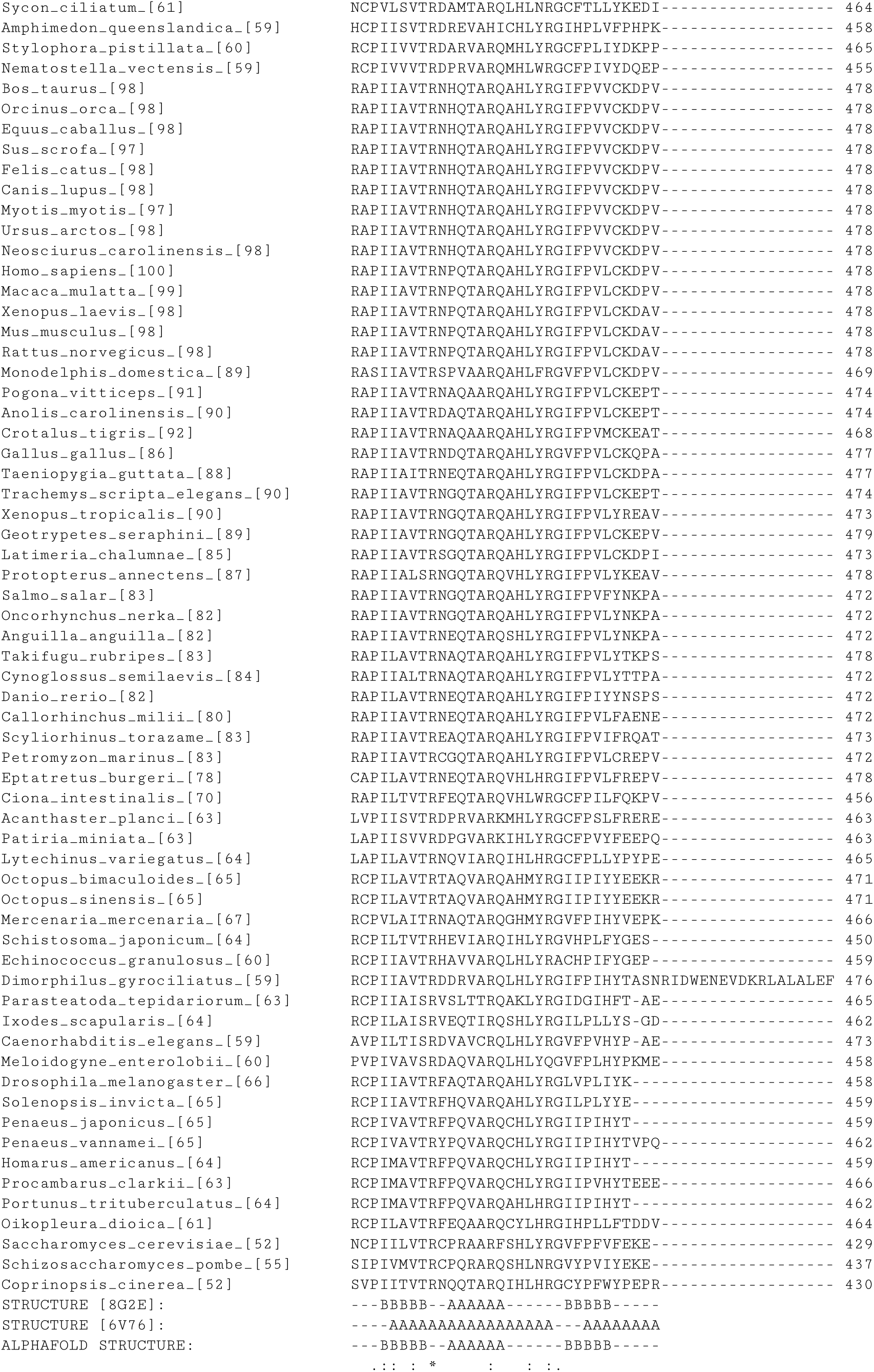
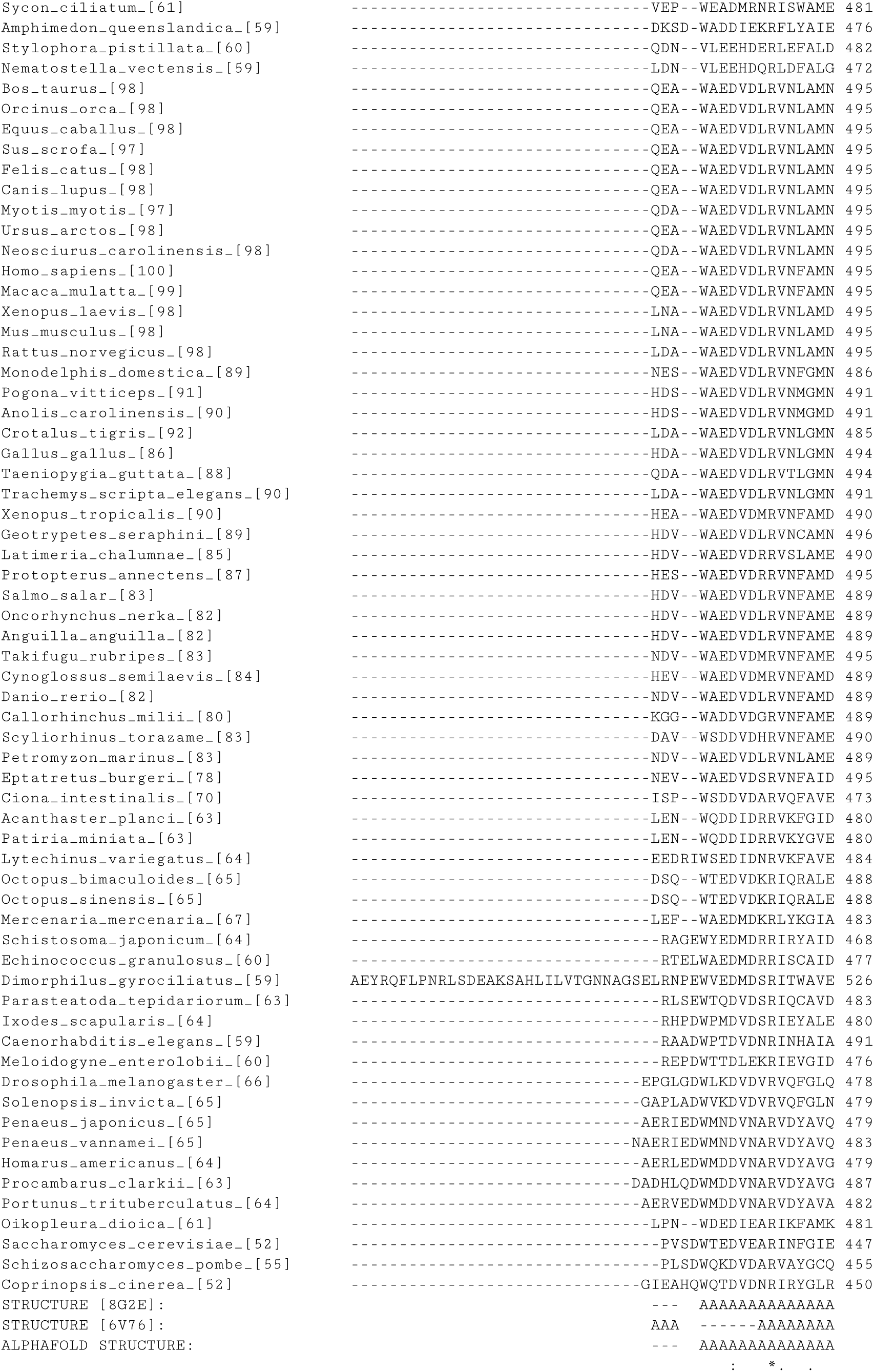
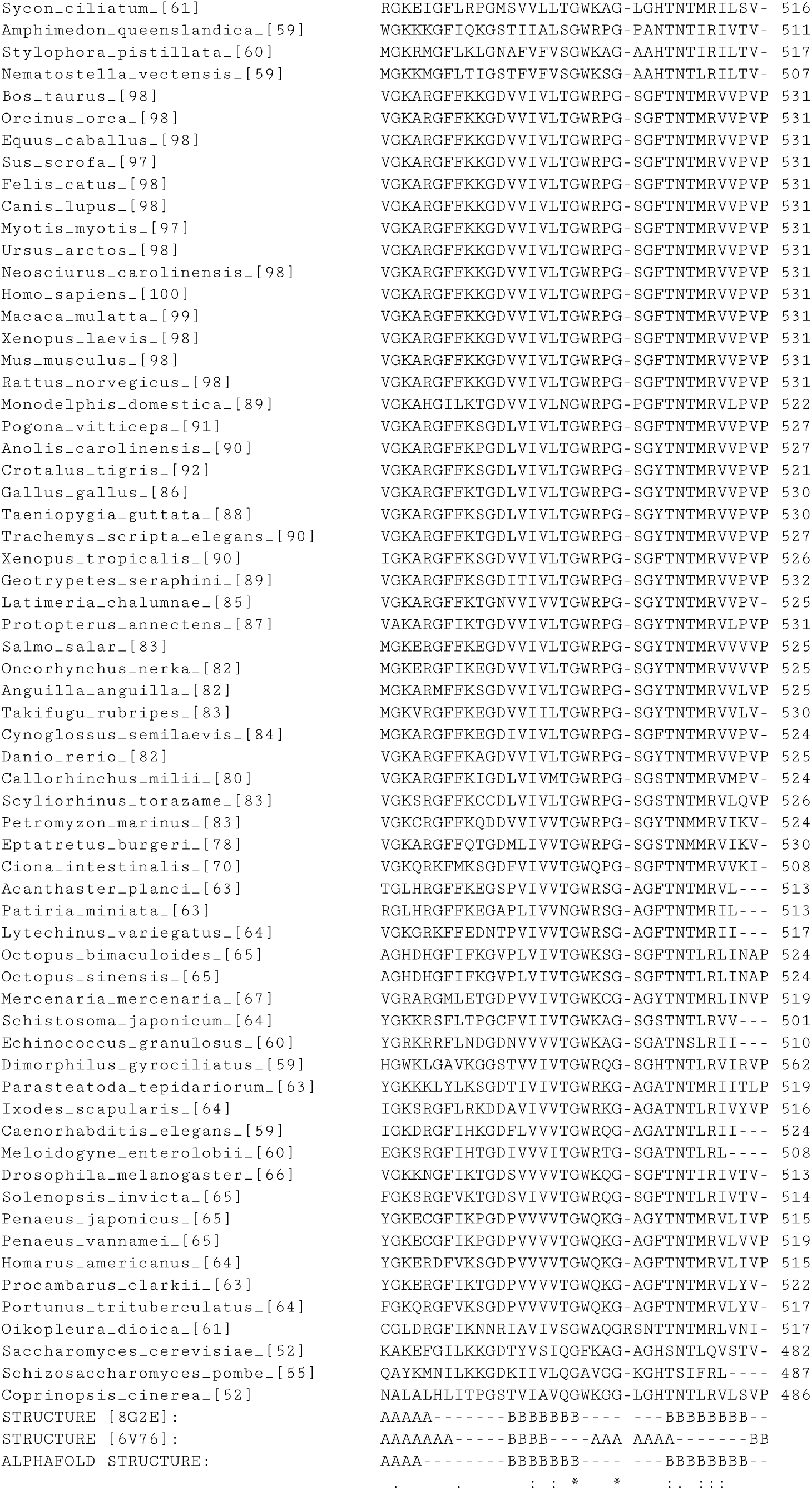
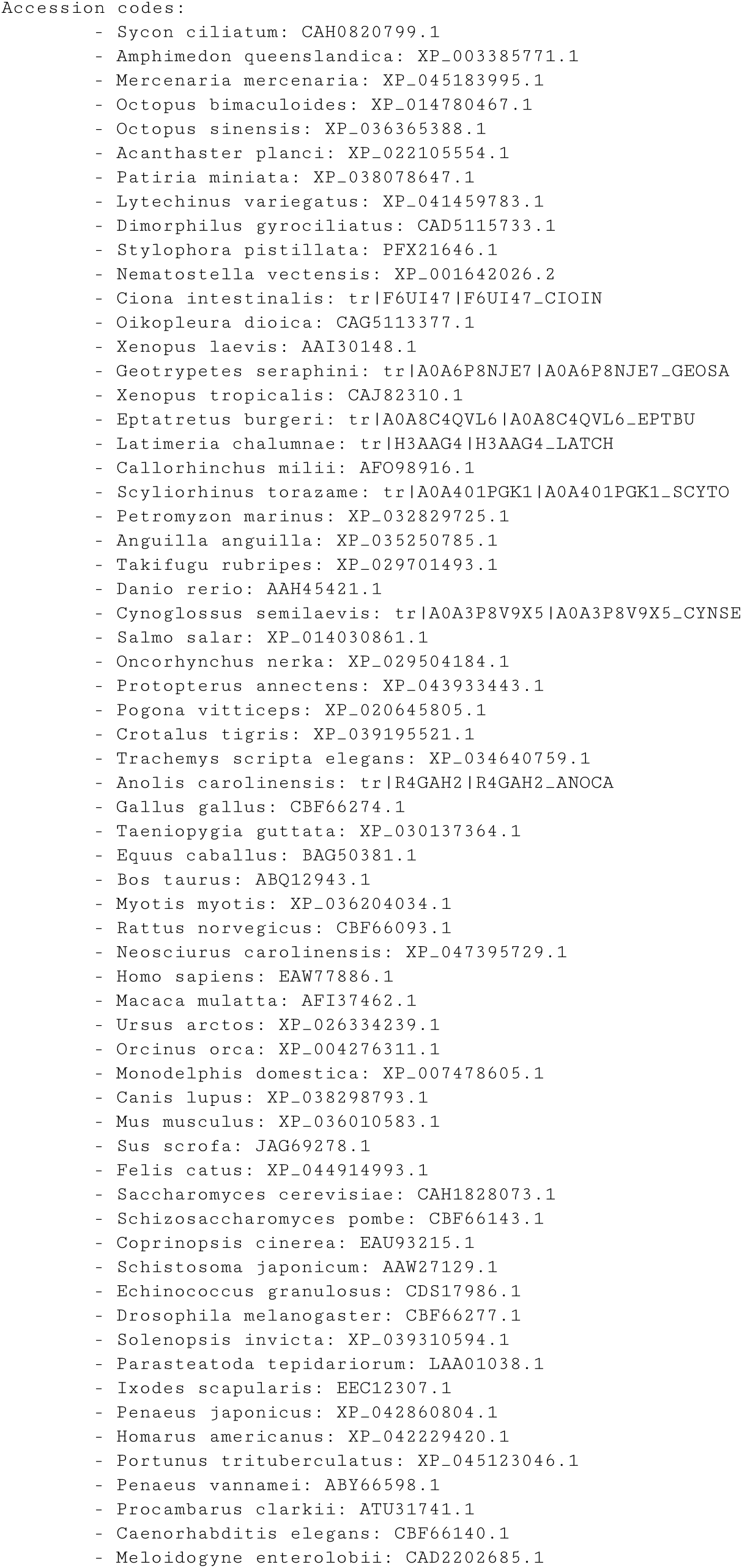
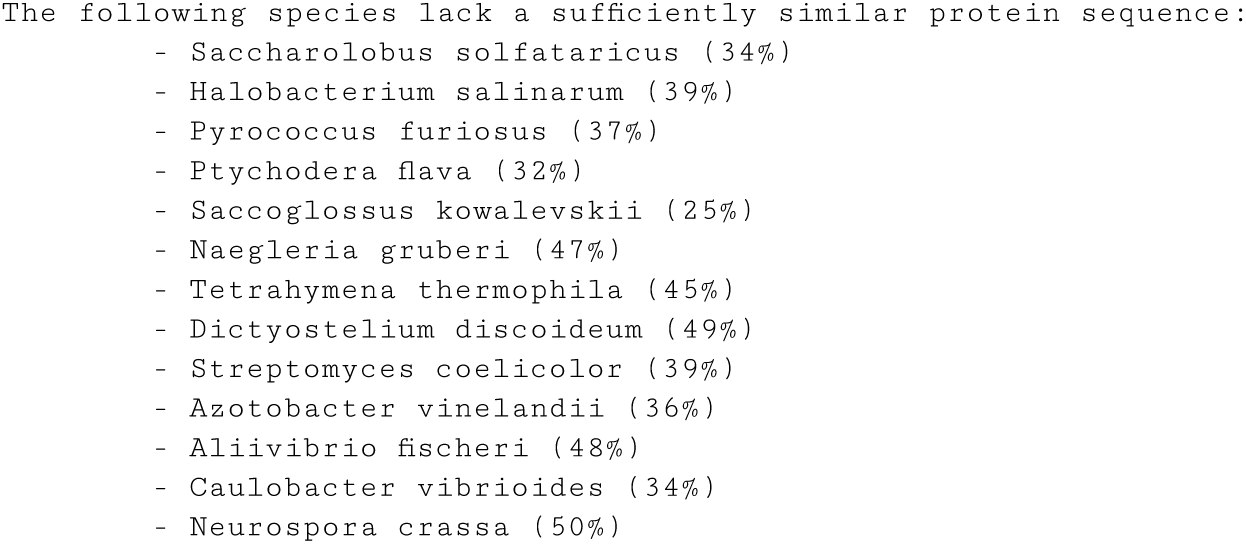
ProteoSync sequence alignment output file from the analysis of human pyruvate kinase (PKM).

